# DSF signalling integrates c-di-GMP and σ⁵⁴ pathways with metabolic reprogramming to control *Stenotrophomonas maltophilia* pathogenicity and antibiotic resistance

**DOI:** 10.64898/2026.02.02.703240

**Authors:** Marc Bravo, Andrómeda-Celeste Gómez, Òscar Conchillo-Solé, Andrea García-Navarro, Joan Lluís Pons, Xavier Daura, Isidre Gibert, Daniel Yero

## Abstract

*Stenotrophomonas maltophilia* is an opportunistic, multidrug-resistant pathogen whose pathogenicity is mainly driven by biofilm formation, extracellular enzyme production, surface adhesins and motility. In addition, a diffusible signal factor (DSF)-mediated quorum sensing (QS) system encoded by the *rpf* gene cluster coordinates collective behaviours and contributes to pathogenicity. While DSF signalling has been extensively studied in plant-pathogenic xanthomonads, its integration with transcriptional and metabolic regulation in human pathogens remains poorly defined. Here, we investigate how the DSF/Rpf system shapes virulence and adaptation in *S. maltophilia* by modulating downstream regulatory and metabolic pathways. Cell density-dependent activation of the RpfC-RpfG two-component system reduces intracellular c-di-GMP levels, promoting a growth phase-dependent switch between biofilm formation and motile lifestyle. This transition requires the global transcriptional regulator Clp, which functions as a transcriptional activator when unbound to c-di-GMP. Clp controls genes involved in motility, adhesion, and the alternative sigma factor σ⁵⁴ RpoN2, which inversely regulates flagellar motility and type IV pilus-mediated adhesion. Transcriptomic profiling uncovered additional DSF-dependent regulatory circuits linking quorum sensing to metabolism and antibiotic resistance. Notably, at the onset of stationary phase, the σ⁵⁴ paralog RpoN1 acts together with RpfF and RpfB to fine-tune DSF production through fatty acid and central carbon metabolism, coupling QS output to cellular metabolic state. This metabolic control constrains DSF output and impacts colistin susceptibility, highlighting clinically relevant consequences of DSF homeostasis. Together, these findings define a species-specific DSF regulatory architecture in *S. maltophilia* that integrates quorum sensing, second-messenger signalling, transcriptional regulation, and metabolic reprogramming to promote survival and pathogenicity.

**Author Summary:** *Stenotrophomonas maltophilia* is a bacterial pathogen that causes infections, particularly in hospitalized and immunocompromised patients. Its clinical impact is driven by intrinsic multidrug resistance and the ability to switch between a free-swimming state and a surface-attached biofilm state. Biofilms protect bacteria from antibiotics and the immune system, making infections persistent, while motility promotes spread and colonization. Understanding how *S. maltophilia* controls this balance is critical for combating infection. Like many bacteria, *S. maltophilia* uses chemical communication, known as quorum sensing, to sense population density and coordinate behaviours linked to virulence and survival. This system is based on the autoinducer DSF, allowing bacteria to decide when to move, attach, or alter metabolism. In this work, we show that DSF-based quorum sensing activates a regulatory cascade that modulates levels of the signalling molecule c-di-GMP, triggering a switch between biofilm formation and motility. This switch relies on global regulators that control genes involved in movement, adhesion, metabolism, and antibiotic resistance. We also uncover a metabolic feedback mechanism that fine-tunes DSF concentration, impacting bacteria state and antibiotic susceptibility. These findings reveal how bacterial communication, metabolism, and pathogenic behaviour are linked in *S. maltophilia*, highlighting opportunities for antimicrobial intervention.

## Introduction

*Stenotrophomonas maltophilia* is an increasing public health concern, with rising infection rates, particularly pneumonia, in recent years [1]. Severe infections can be life-threatening in immunocompromised patients, those in intensive care units, and individuals with cystic fibrosis [2–4]. Clinical management is challenging due to molecular heterogeneity, intrinsic multidrug resistance and capacity to form biofilms on medical devices [5,6]. Pathogenesis relies on diverse virulence factors, including surface polysaccharides and adhesins, flagella, type 1 fimbriae (SMF-1), type IV pili, lytic enzymes, proteases, and iron acquisition systems [4,7]. In addition, the diffusible signal factor-mediated quorum sensing (DSF-QS) system regulates metabolism and virulence gene expression, promoting colonization, biofilm formation, and interspecies interactions [2,4,7,8]. Consistent with the molecular heterogeneity, two distinct DSF-QS genetic variants have been identified among S. maltophilia strains, each associated with differences in virulence-related traits and antibiotic resistance profiles [9–11].

The DSF molecule produced by *S. maltophilia* is cis-11-methyl-2-dodecenoic acid, synthesized by RpfF, an enoyl-CoA hydratase encoded in the *rpf* (regulation of pathogenicity factors) gene cluster [8,9,12]. The DSF-QS system and *rpf* genes were first identified in *Xanthomonas campestris pv. campestris* (Xcc) and other *Lysobacteraceae* (≡ *Xanthomonadaceae*) [13]. In Xcc, the RpfC-RpfG two-component system regulates QS through signal perception and transcriptional control, while RpfB, a long-chain fatty acyl-CoA ligase, participates in DSF turnover [14–16]. RpfC is a hybrid sensor with transmembrane, histidine kinase (HK), CheY-like receiver (REC), and histidine phosphotransferase (HPT) domains. At low cell density, the REC domain binds RpfF, inhibiting DSF production; at higher densities, DSF sensing triggers REC phosphorylation and RpfF release, completing DSF synthesis [11,17,18]. RpfG, the cognate response regulator, receives the phosphoryl group from RpfC and activates its phosphodiesterase activity, degrading intracellular cyclic dimeric GMP (c-di-GMP) [19,20]. In *Xcc*, c-di-GMP binds the global regulator Clp (cAMP receptor-like protein), controlling transcription, metabolism, biofilm formation, and motility [20–22]. This DSF/Rpf-Clp axis forms a hierarchical network modulating multiple biological functions. In *Xcc* DSF synthesis is further influenced by lipid metabolism under the control of the σ⁵⁴ alternative sigma factor RpoN1 [23], adding complexity to the system

The DSF/Rpf signalling system of *S. maltophilia* is relatively conserved with respect to its homologous system in *Xanthomonas sp*. (Fig 1A). The roles of RpfF and RpfC in DSF synthesis and sensing have been investigated through mutagenesis, and experimental evidence further confirms that an RpfF-RpfC complex regulates DSF production [9,11]. On the other hand, a systematic analysis of c-di-GMP turnover enzymes with diguanylate cyclase or phosphodiesterase activity revealed that RpfG regulates swimming motility and biofilm formation in *S. maltophilia* [24]. Furthermore, the global regulator Clp in *S. maltophilia* has been implicated in temperature-dependent protease production and biofilm formation through the LotS-LotR and RpfC-RpfG two-component systems, with c-di-GMP acting as a negative effector of their activity [25,26]. Despite significant advances in understanding the DSF-QS system in *S. maltophilia*, the precise molecular mechanisms controlling DSF-mediated regulation remain unclear, and the full spectrum of virulence factors activated by this system is not yet known.

**Fig 1.**
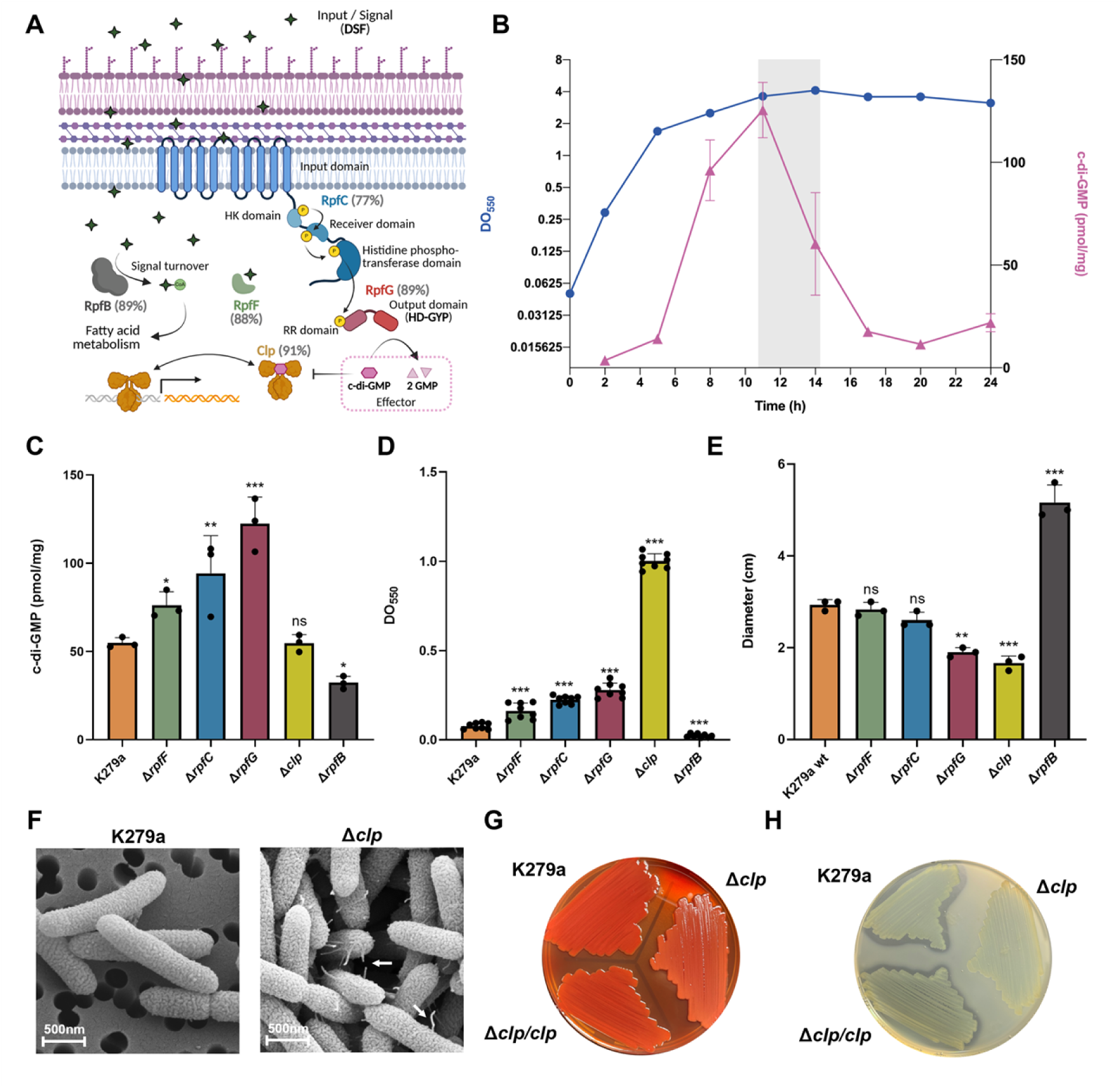
DSF/Rpf–c-di-GMP–Clp signalling controls the transition between planktonic and biofilm lifestyles in *S. maltophilia* K279a. (A) Schematic model of the DSF quorum-sensing (QS) pathway in *S. maltophilia* K279a based on homology with *Xanthomonas campestris* pv. *campestris* (*Xcc*). Sequence identity with the *Xcc* ortholog is indicated in parentheses. DSF is synthesized by RpfF and its turnover involves RpfB; the hybrid sensor RpfC perceives DSF and phosphorylates the phosphodiesterase RpfG, which degrades c-di-GMP. The global regulator Clp mediates downstream transcriptional responses. (B) Time course of bacterial growth (blue circles, OD₅₅₀) and intracellular c-di-GMP concentration (pink triangles, pmol mg⁻¹ protein) in LB at 37 °C. The shaded area denotes the previously identified time window of maximal DSF production (Coves et al., 2023). Means ± SEM of three independent cultures are shown. (C) Intracellular c-di-GMP levels after 14 h of growth, quantified by ELISA (pmol mg^-1^ protein). (D) Biofilm biomass quantified by crystal violet staining (OD_550_) after 24 h in BM2 at 37 °C. (E) Swimming motility assessed on LB with 0.25 % agar at 30 °C for 48 h; bars show colony diameter. In panels C–E, values represent means ± SD (n ≥ 3); significance was evaluated by one-way ANOVA with Tukey’s post hoc test (*, *p* < 0.05; **, *p* < 0.01; ***, *p* < 0.001; ns, not significant). (F) Scanning electron micrographs of wild-type and Δ*clp* cells reveal abundant pili/fimbriae in the Δ*clp* mutant (white arrows) absent from wild type. Scale bars, 500 nm. (G) Qualitative analysis of extracellular polysaccharide (EPS) production on BHI agar containing 3% sucrose and Congo red. (H) Exoproteolytic activity visualized as clearing on LB–5% egg yolk agar.

In this study, we aim to define the DSF regulon in *S. maltophilia* and its connection with pathogenic phenotypes by investigating the interplay between DSF synthesis and signalling, intracellular c-di-GMP levels and transcriptional regulators including sigma factor σ⁵⁴. By resolving DSF-dependent regulation across growth phases, we assess whether DSF signalling in *S. maltophilia* operates through a regulatory architecture that diverges from that described in plant-pathogenic xanthomonads. We explore whether c-di-GMP functions as a molecular switch that governs the lifestyle transition from motility to biofilm in *S. maltophilia*, as observed in related bacterial pathogens [27]. In this context, we uncover a direct Clp-σ⁵⁴ regulatory axis that links quorum sensing to motility-biofilm switching and metabolic reprogramming. We further elucidate the role of DSF in connecting core metabolic pathways, particularly fatty acid (FA) metabolism, to stress adaptation and the regulation of virulence traits, including biofilm formation and colistin resistance, thereby repositioning DSF signalling as a species-adapted regulatory system with clinically relevant consequences in a multidrug-resistant human pathogen.

## Results

### The DSF/Rpf-c-di-GMP-Clp signalling cascade controls the switch between the planktonic and biofilm modes of growth

Variation in c-di-GMP levels in *S. maltophilia* K279a was first assessed by quantifying this second messenger throughout bacterial growth (Fig 1B). Cells were grown in LB medium at 37 °C with agitation in triplicate, and samples were collected over 24 h. Intracellular c-di-GMP levels gradually increased during exponential growth, reaching a maximum at 11 h, before declining more rapidly as cells prepared to enter stationary phase. This behaviour coincided with the peak of DSF production upon entry into stationary phase (11-14 h) (Fig 1B), as we had previously shown [28]. To examine the link between declining intracellular c-di-GMP levels and activation of the DSF quorum-sensing system in *S. maltophilia*, we generated and analysed unmarked deletion mutants in K279a targeting key QS-associated genes involved in DSF synthesis (*rpfF*; *smlt2235*) and sensing (*rpfC*; *smlt2234*), signal transduction (*rpfG*; *smlt2233*), putative DSF turnover (*rpfB*; *smlt2236*), and the global transcriptional regulator Clp (*smlt4306*).

Intracellular c-di-GMP levels were measured at stationary phase (14 h), when DSF-QS is active (Fig 1C). Deletion of *rpfF*, *rpfC*, or *rpfG* significantly increased c-di-GMP, with Δ*rpfG* showing >2-fold accumulation, consistent with loss of RpfG phosphodiesterase activity. In contrast, *ΔrpfB* had reduced c-di-GMP, supporting its role in DSF turnover and RpfG activation. Deletion of *clp* had no detectable effect on c-di-GMP concentration. Complemented strains restored or further reduced c-di-GMP levels (S1 Fig), confirming gene-specific effects. DSF production analysis in mutants confirmed the role of RpfF in autoinducer synthesis and RpfC-mediated repression of DSF via RpfF sequestration [9,29]. Elevated DSF in *ΔrpfB* supports its function as a DSF-degrading enzyme, whereas *ΔrpfG* or *Δclp* had no effect on DSF levels (S1 Fig).

Phenotypic analyses showed that deletion of *rpfF*, *rpfC*, *rpfG* or *clp* significantly increased biofilm formation compared with the wild-type (Fig 1D). In contrast, swimming motility was significantly reduced only in the Δ*rpfG* and Δ*clp* mutants (Fig 1E). *Δclp* also displayed more surface pili and fimbriae (Fig 1F), decreased EPS (Fig 1G), and loss of exoproteolytic activity (Fig 1H). Conversely, *ΔrpfB* exhibited reduced biofilm and increased motility (Fig 1D and E). All phenotypes were restored by complementation (Fig 1G, H and S1). All results are consistent with c-di-GMP levels in these mutants and the known link between high c-di-GMP and biofilm formation. These results also indicate that DSF-QS activation mediates biofilm dispersion via the DSF/Rpf-c-di-GMP-Clp cascade. Additionally, *ΔrpfF* showed increased and *ΔrpfB* reduced susceptibility to colistin, with no changes for six other antibiotic classes (S1 Table).

### DSF-QS-mediated genetic regulatory network in *S. maltophilia*

Transcriptomes of *rpf* cluster mutants and the *clp* mutant were analysed at maximal DSF production (14 h of growth in LB at 37 °C) (Fig 2 and S1 File). At this stage, mutant strains displayed differences in the number and direction of differentially expressed genes (DEGs), with the strongest transcriptional impact observed in Δ*clp* compared with the other mutants (Fig 2A-C). In mutants Δ*rpfF*, Δ*rpfC*, Δ*rpfG* and Δ*clp*, 60-80% of DEGs were downregulated, indicating that DSF-QS predominantly activates gene expression. Consistent with its distinct phenotype, the Δ*rpfB* mutant showed an inverse transcriptional profile, with 83% of DEGs upregulated (Fig 2A-C). COG analysis showed broad regulation of biological processes for all mutants with an opposite pattern of downregulated and upregulated genes in each functional category for mutant Δ*rpfB* compared to the other four mutants that are part of the DSF/Rpf-Clp axis (Fig 2D). Sixteen DEGs were shared among these four mutants, 12 of which were consistently downregulated (Table 1, Fig 2B). Only a tRNA-Leu gene and the pilin genes *pilA* (*smlt3757*) and *pilE* (*smlt3758*) were commonly upregulated, whose increased expression likely underlies the enhanced biofilm formation observed in these mutants through elevated type IV pili production [30]. Δ*rpfC*, Δ*rpfG* and Δ*clp* mutants shared 92 DEGs (Fig 2A-C and S1 File), with the most highly represented COG categories associated with envelope biogenesis and motility, both negatively regulated (Fig 2D). Interestingly, 6-15% of DEGs were mutant-specific (Fig 2B and C), suggesting additional DSF-QS regulatory routes independent of the DSF/Rpf-Clp axis.

**Fig 2.**
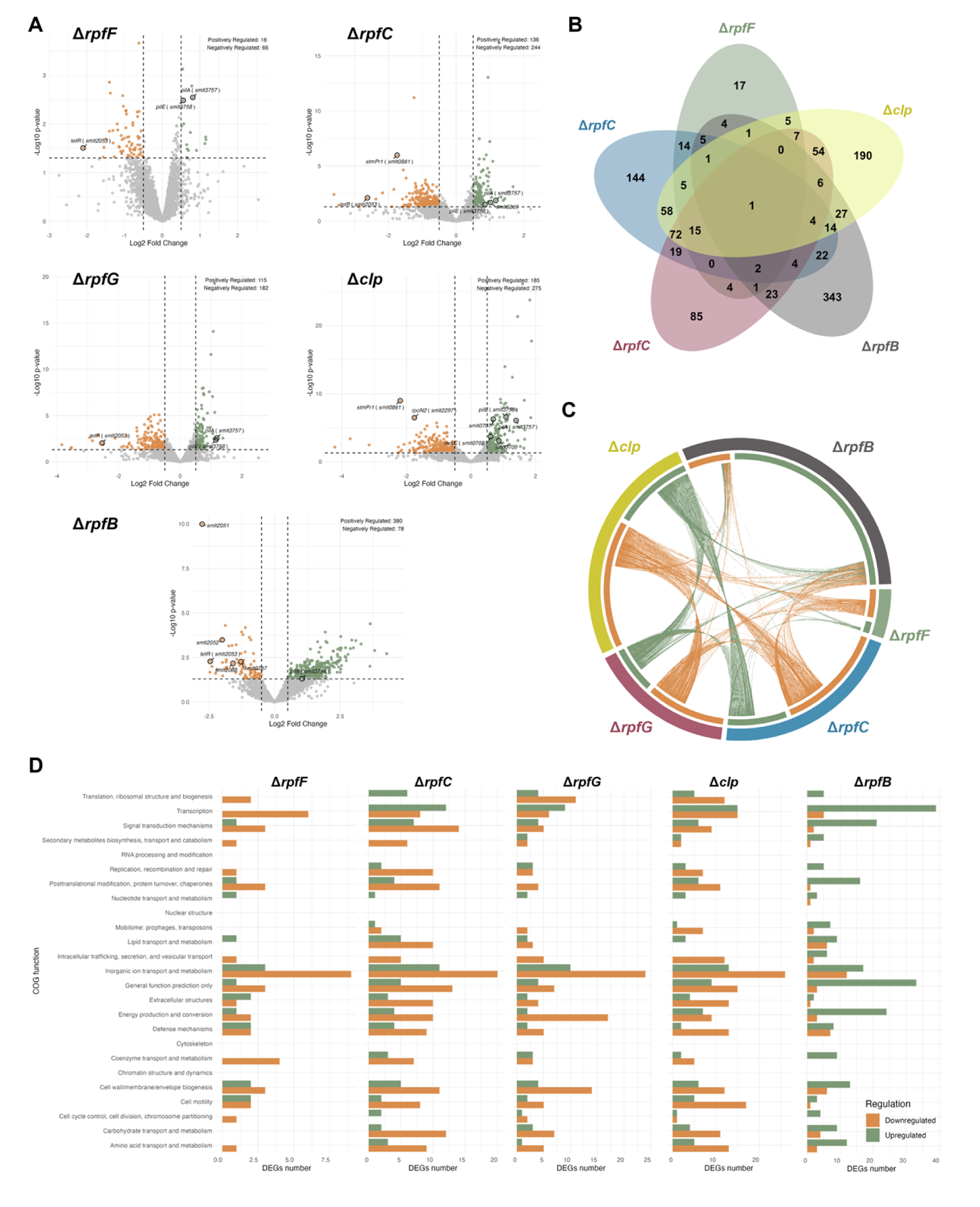
Transcriptomic profiling of the DSF/Rpf–Clp signalling cascade in *S. maltophilia* K279a. (A) Volcano plots showing differentially expressed genes (DEGs) in Δ*rpfF*, Δ*rpfC*, Δ*rpfG*, Δ*clp*, and Δ*rpfB* mutants compared with the wild type at 14 h (DSF quorum-sensing activation). Genes with |log₂ fold change| ≥ 0.5 and P < 0.05 are highlighted (green, upregulated; orange, downregulated). Selected DEGs are labelled, including *pilA* (*smlt3757*), *pilB* (*smlt3756*), *pilE* (*smlt3758*), *rpoN2* (*smlt2297*), *fleQ* (*smlt2295*), *stmPr1* (*smlt0861*), the type 1 fimbriae operon *smlt0706–0709* (SMF-1), and the β-oxidation operons *smlt2053–2051* and *smlt0264–0268*, shown only when deregulated. (B) Venn diagram depicting the overlap of DEGs among Δ*rpfF*, Δ*rpfC*, Δ*rpfG*, Δ*clp*, and Δ*rpfB* mutants, illustrating shared and unique transcriptional responses. (C) Circos plot representing DEGs common to two or more mutants, highlighting transcriptional convergence and divergence within the DSF/Rpf–Clp axis. Green lines denote downregulated and orange lines upregulated genes. (D) COG functional classification of DEGs in each mutant at 14 h. Bars represent the number of downregulated (green) or upregulated (orange) genes in each functional category.

**Table 1.**
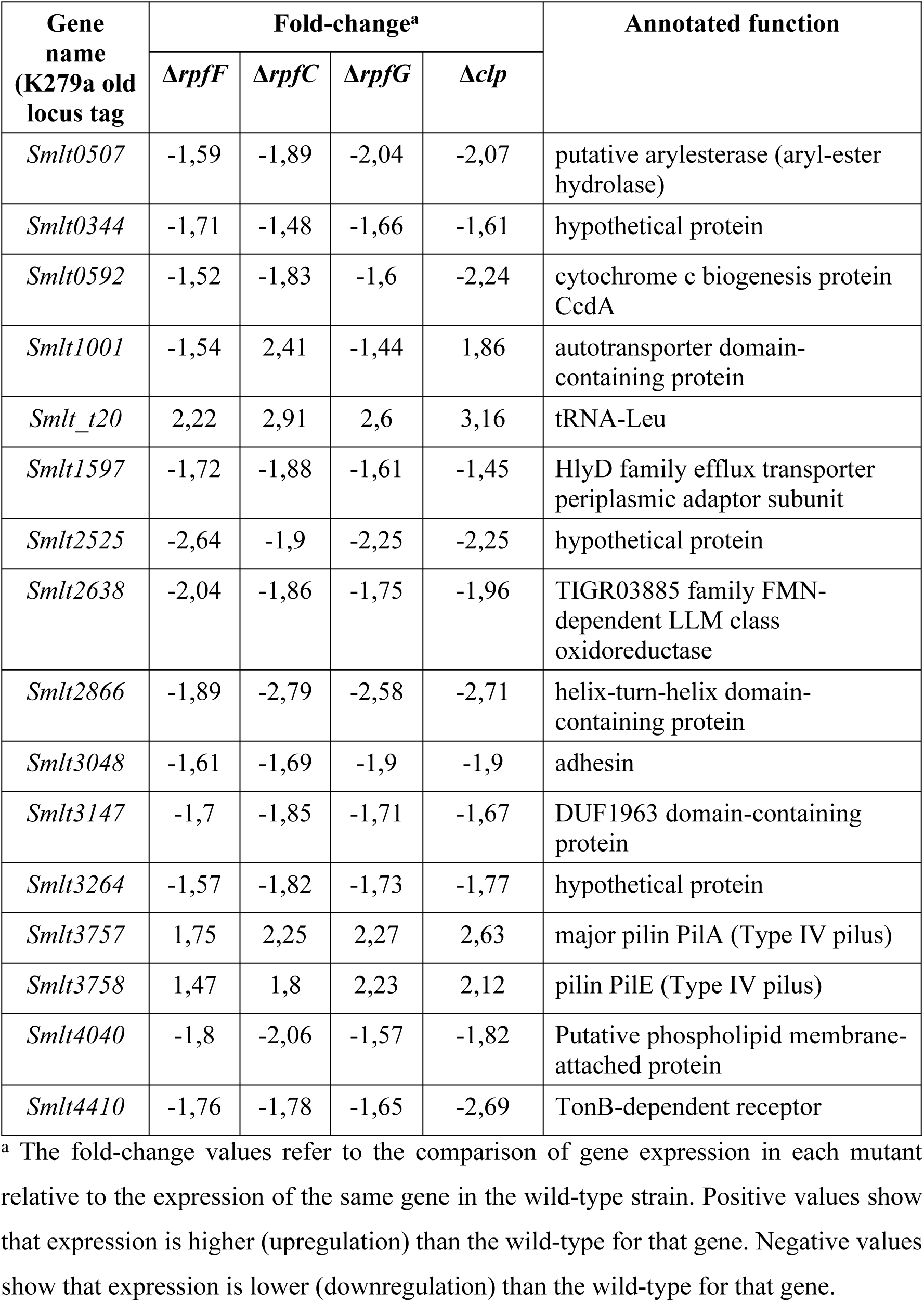
Common differentially expressed genes (DEG with *p*-value<0.05) amongst four *S. maltophilia* K279a mutant strains lacking individual components of the DSF/Rpf-Clp signalling pathway.

To control for growth phase-dependent effects and to depict a broader regulatory network, global transcriptional changes in wild-type K279a between 5 and 14 h under identical growth conditions were analysed (S2 Fig and S1 File). This comparison identified 3,076 DEGs (68% of the genome), with similar numbers of down- and upregulated genes. At stationary phase entry, most rpf cluster genes were modestly downregulated, whereas *rpfF* was upregulated 2.66-fold (S2 Fig), consistent with increased DSF production at high cell density. Functional over-representation analyses (COG/GO/KEGG) showed that upregulated DEGs were associated with protein synthesis, ribosome biogenesis and inorganic ion transport and metabolism, with KEGG highlighting ribosome, aminoacyl-tRNA biosynthesis and protein export pathways. Iron acquisition operons and FA metabolism were also enriched, consistent with DSF as a FA-derived signal. In contrast, downregulated DEGs were enriched for regulatory functions, with over-representation of the COG categories transcription and signal transduction, together with repression of pili-associated operons and pathways connected to nitrogen metabolism and amino-acid degradation. Among the global transcriptional regulators that are significantly deregulated at the onset of the stationary phase, a metalloregulator of the ArsR/SmtB family (*smlt2419*; −7.94-fold), the two-component system response regulator RegA (*smlt1784*; −4.77-fold), and a σ⁵⁴-like sigma factor (*smlt1112*; −1.69-fold) stand out (S2 Fig and S1 File). Comparison with mutant transcriptomes revealed that 78-82% of DEGs overlapped with those identified in the wild-type at 14 h (S2 Fig), indicating that the Rpf-Clp pathway primarily modulates transcriptional programs intrinsic to the transition into stationary phase rather than inducing an entirely distinct regulon.

### Growth phase-dependent activation of the Clp regulon links DSF-QS to the planktonic-biofilm transition

*S. maltophilia* activates DSF-QS system at the onset of stationary phase, likely modulating gene expression via the c-di-GMP-dependent regulator Clp. To examine this mechanism, we analysed the transcriptome of the K279a Δ*clp* mutant at late exponential growth (5 h), when c-di-GMP begins to accumulate, and at stationary phase entry (14 h), when DSF peaks and c-di-GMP start to decline. In the wild-type at 5 h, discrete levels of c-di-GMP in the absence of DSF is expected to inhibit Clp activity, whereas maximal RpfG phosphodiesterase activity at 14 h (DSF-QS activation) should promote c-di-GMP degradation and progressive Clp-dependent gene activation.

During exponential growth, the K279a Δ*clp* mutant displayed 305 DEGs, the majority of which were upregulated (66%) (Fig 3A). Genes associated with flagellar assembly, motility, and chemotaxis and signal transduction mechanisms were significantly upregulated (Fig 3C, D, and S3), contrasting with the reported role of Clp as a transcriptional activator at low c-di-GMP levels [31] and suggesting indirect regulation via downstream factors in the exponential growth phase. Notably, downregulated genes included the type IV pilins *pilA* and *pilE*, as well as a second σ⁵⁴-like sigma factor (*smlt2297*) and the σ⁵⁴-dependent activator FleQ (*smlt2295*) (Fig 3A, C and E). As σ⁵⁴ factor and FleQ form a hierarchical regulatory module controlling flagellar gene expression in *Legionella pneumophila* [32], these regulators are likely directly controlled by Clp in *S. maltophilia* and they mediate broader transcriptional effects in a hierarchical manner.

**Fig 3.**
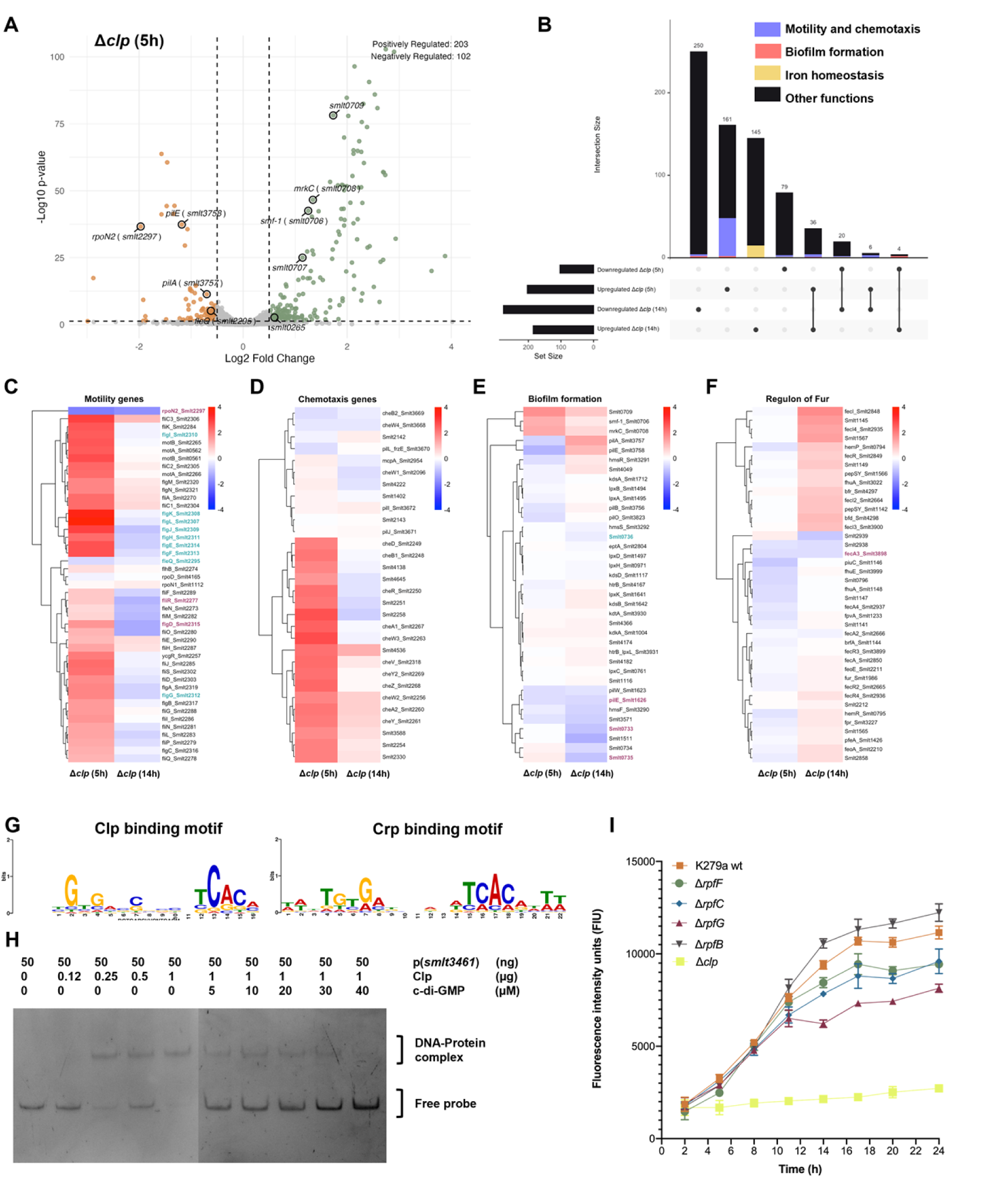
Growth phase–dependent activation of the Clp regulon connects DSF signalling with the planktonic–biofilm transition in *S. maltophilia* K279a. (A) Volcano plot of differentially expressed genes (DEGs) in the Δ*clp* mutant relative to the wild type at mid–late exponential phase (5 h). Genes with |log₂ fold change| ≥ 0.5 and *p* < 0.05 are highlighted (green, upregulated; orange, downregulated). The same selected DEGs as in Figure 2 are labelled, shown only when deregulated. (B) UpSet plot showing the overlap of DEGs in Δ*clp* mutants at 5 h and 14 h, highlighting functional categories associated with motility and chemotaxis (blue), biofilm formation (red), and iron homeostasis (yellow). (C–F) Heat maps of DEGs involved in motility (C), chemotaxis (D), biofilm formation (E), and iron homeostasis (regulon of Fur) (F) comparing Δ*clp* cells at 5 h and 14 h with wild-type controls at the same time points. Red indicates upregulation and blue downregulation. Genes whose promoter regions contain the conserved Clp-binding motif are highlighted in purple; genes in blue lack the motif but belong to operons containing red-labelled genes. (G) Conserved palindromic motif (left) identified by MEME within promoters of downregulated DEGs in the Δ*clp* mutant at 14 h (consensus TGTGA-N₆-TCACA) and its closest match, the Gammaproteobacteria Crp binding motif (right) identified with TomTom. (H) Electrophoretic mobility shift assay (EMSA) demonstrating Clp binding to the promoter of *smlt3461* in vitro. Increasing Clp concentrations (left gel) shifted the probe from the free form to a DNA–protein complex, whereas addition of c-di-GMP (right gel) progressively inhibited complex formation. (I) Fluorescence intensity over time in K279a wild-type and indicated mutants carrying a chromosomal Psmlt3461-sfGFP reporter. Data show Clp-dependent activation of *smlt3461* inversely correlated with intracellular c-di-GMP levels.

As shown above, at stationary phase entry, the K279a Δ*clp* mutant displayed an expanded transcriptional response, with 185 genes upregulated (40%) and 275 downregulated (60%) (Fig 2A), 82% of which overlapped with wild-type DEGs at this stage (S3 Fig). Functional enrichment indicated that Clp acts primarily as a positive regulator, affecting intracellular trafficking and secretion, inorganic ion transport, extracellular structures, defence mechanisms, carbohydrate metabolism and motility (Fig 2D). Adhesion- and motility-related genes showed growth phase-dependent regulation. Chemotaxis and flagellar genes were largely downregulated, whereas adhesion-associated genes, including *pilA*, *pilB* (*smlt3756*), *pilE* and the SMF-1 fimbrial operon (*smlt0706-0709*), were upregulated (Fig 3B-E). Motility genes were inversely regulated in the Δ*clp* mutant relative to the wild-type at stationary phase entry (Fig 3C and S3), supporting a role for Clp in DSF-QS-dependent motility control. As in exponential phase, the σ⁵⁴ factor Smlt2297 and the σ⁵⁴-dependent activator FleQ remained downregulated (Fig 3C).

In addition to pilins and fimbriae, other differentially expressed virulence genes help explain the K279a Δ*clp* mutant phenotypes observed in Fig 1 (see S1 File). At stationary phase entry, the major extracellular protease gene StmPr1 (*smlt0861*) was strongly downregulated (−4.57-fold). At this stage (DSF-QS activation), most Fur regulon genes were significantly upregulated in the Δ*clp* mutant (Fig 3F), but not during exponential growth, indicating that Clp affects iron homeostasis genes in a growth-dependent and it likely occurs indirectly via another regulator.

### Clp activates target gene expression by binding to their promoter regions in the absence of c-di-GMP

To explore Clp-mediated regulation, we used MEME Suite to identify conserved motifs in promoters of DEGs in the K279a Δ*clp* mutant. At DSF-QS activation, a palindromic motif TGTGA-N₆-TCACA was found in 180 of 275 downregulated DEGs (*p*-value < 5 × 10⁻³) (Fig 3G and S1 File). TomTom analysis showed the motif matched the Crp binding motif in Gammaproteobacteria (*p*-value = 5.28 × 10⁻⁶ and *q*-value = 2.67 × 10⁻³), notable because Crp is considered a homologue to Clp [31]. This motif is also the consensus Clp binding site in *Xcc*, where Clp regulates gene expression via promoter binding [33]. No conserved motif was found in most DEGs at 5 h or in upregulated genes at 14 h, suggesting regulation may occur indirectly. RNA-seq from the Δ*clp* mutant revealed deregulation of transcription factors including σ⁵⁴ factor Smlt2297, LysR family transcriptional regulator Smlt2509 (−2.81-fold), CopG family transcriptional repressor Smlt1292 (−2.41-fold), and FtrA family transcriptional regulator Smlt2098 (−2.21-fold); all contained the Clp binding motif in their promoters (S1 File). However, only a few genes related to motility, biofilm formation, and Fur regulation carried this motif (Fig 3C-F).

To confirm Clp binding to the motif TGTGA-N₆-TCACA, we purified Clp and performed EMSA using a DNA fragment from the *smlt3461* promoter, which contains a nearly identical site (TGTGA-N₆-TCACC) and is downregulated in the Δ*clp* mutant at both time points (−2.97 and −1.44-fold). EMSA showed progressive DNA-protein complex formation as the concentration of Clp increases, reaching complete binding at 1 μg (Fig 3H). Addition of c-di-GMP (up to 40 μM) gradually released the DNA, fully inhibiting binding at the highest concentration, consistent with prior evidence of high-affinity for the c-di-GMP-Clp interaction (Kd ≈ 2 × 10⁻⁷ M) [34]. These results indicate that Clp binds the *smlt3461* promoter in the absence of c-di-GMP, and binding is reversibly inhibited by c-di-GMP *in vitro*.

To validate these findings *in vivo*, c-di-GMP-sensing *S. maltophilia* reporter strains with different genetic backgrounds (K279a mutants and the wild-type) were constructed in which the gene for GFP was inserted chromosomally and under the control of the promoter region of the *smlt3461* gene (see methods). Fluorescence was measured at the same time points as c-di-GMP quantification in Fig 1B. In the Δ*clp* mutant, GFP remained at basal levels, consistent with loss of Clp-mediated activation, while other strains showed increasing fluorescence over time (Fig 3I). Fluorescence inversely correlated with intracellular c-di-GMP levels, with mutants accumulating more c-di-GMP (see Fig 1C) exhibiting lower GFP, most notably the Δ*rpfG* mutant. These results confirm that Clp activates transcription via the TGTGA-N₆-TCACA motif, and this activation is modulated by c-di-GMP and regulated by the DSF-QS system.

### Clp regulates gene expression through direct regulation of the alternative sigma factor σ⁵⁴ RpoN2

Among the global regulators controlled by Clp, the alternative sigma factor σ⁵⁴ (Smlt2297) is notable, being significantly downregulated in the Δ*clp* mutant at both growth phases, suggesting a downstream role in the DSF-QS system. Its *Xcc* ortholog, RpoN2, regulates flagellar biosynthesis, swimming motility, biofilm formation, EPS production, and virulence [35,36] and in S. maltophilia it is proposed at the top of the flagellar cascade [37]. In K279a, this σ⁵⁴ factor forms a putative operon with a two-component system response regulator (*smlt2296*) and *fleQ*, and its promoter contains a palindromic sequence resembling the Clp-binding motif (TGTGA-N₆-TCACA, Fig 4A). EMSA with purified Clp showed concentration-dependent binding to this promoter, with complete probe shift at 1 μg (Fig 4B). Addition of c-di-GMP gradually released the DNA probe, restoring normal migration, confirming c-di-GMP-dependent inhibition of Clp-DNA binding (Fig 4B).

**Fig 4.**
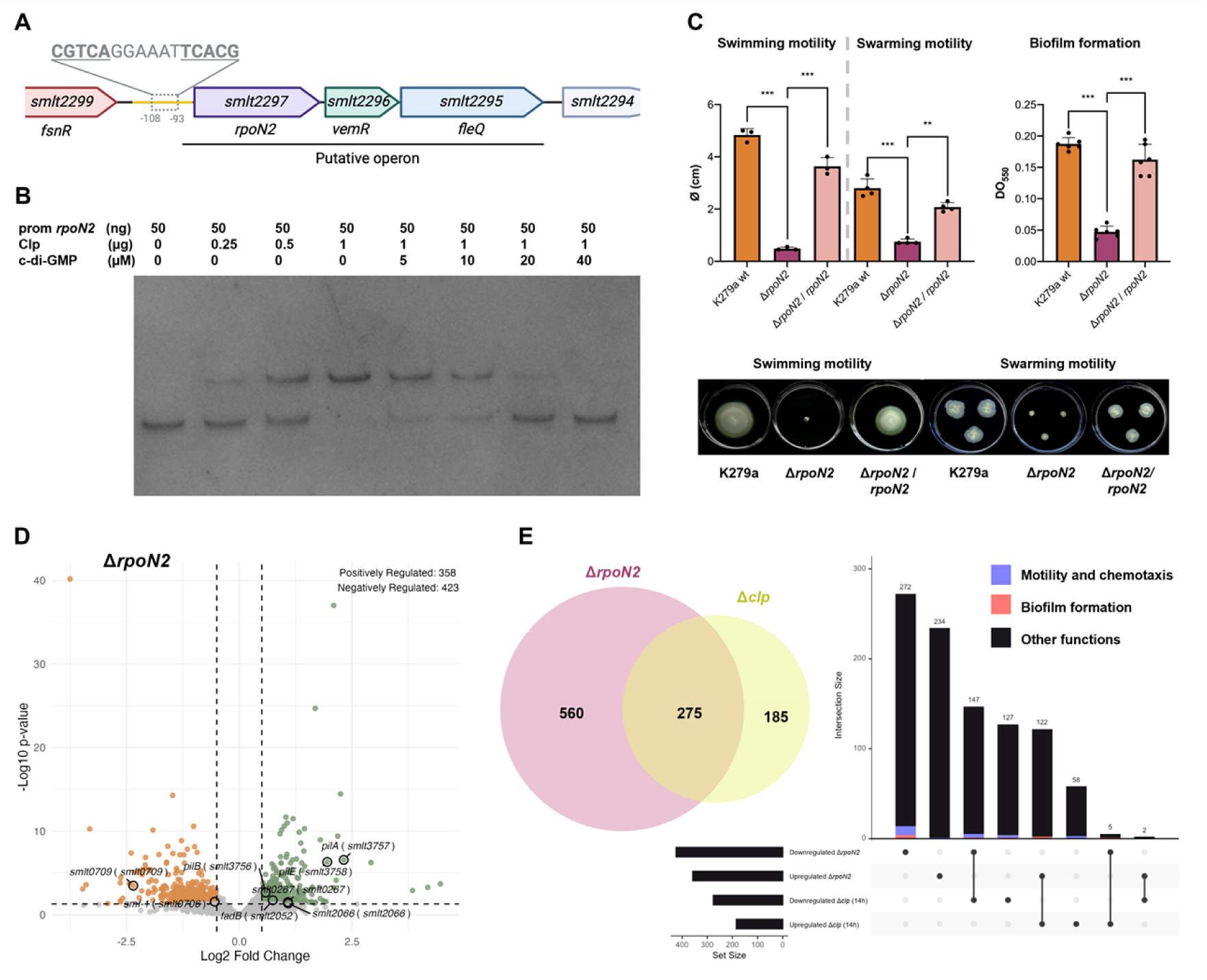
Clp directly activates the σ⁵⁴ factor RpoN2 operon to control motility and biofilm formation in *S. maltophilia* K279a. (A) Schematic representation of the putative *rpoN2* operon (*smlt2297–2294*), which encodes the σ⁵⁴ factor, a response regulator 92% identical to VemR in *Xcc* and the σ⁵⁴-dependent activator FleQ. The upstream promoter harbors a palindromic sequence (highlighted) closely matching the Clp-binding motif (TGTGA-N₆-TCACA). The yellow box marks the DNA fragment amplified for EMSA. (B) Electrophoretic mobility shift assay (EMSA) showing Clp binding to the *rpoN2* promoter. Increasing Clp concentrations progressively shift the probe to a DNA–protein complex, while rising c-di-GMP concentrations disrupt this interaction and restore free-probe migration. (C) Phenotypic analysis of motility and biofilm formation. Bar graphs show swimming and swarming diameters and crystal-violet–quantified biofilm biomass for wild type, Δr*poN2*, and complemented strains after 48 h (swimming), 5 days (swarming), or 24 h (biofilm). Representative assay images are shown below. Bars represent means ± SD (n ≥ 3). Statistical significance was assessed by one-way ANOVA with Tukey’s test (**, *p* < 0.01; ***, *p* < 0.001). (D) Volcano plot of differentially expressed genes (DEGs) in the Δ*rpoN2* mutant compared to wild type at 14 h (DSF quorum-sensing activation). Genes with |log₂ fold change| ≥ 0.5 and p < 0.05 are coloured green (upregulated) or orange (downregulated). The same selected DEGs as in Figure 2 are labelled, shown only when deregulated. (E) Regulatory overlap between Clp and RpoN2. Venn diagram depicts DEGs shared between Δ*rpoN2* and Δ*clp* mutants at 14 h. The accompanying UpSet plot details intersecting DEG sets, highlighting functions linked to motility and chemotaxis (blue) and biofilm formation (red).

To examine Clp regulation of motility via RpoN2, we generated a Δ*rpoN2* mutant in K279a. The mutant lost swimming and swarming motility and showed reduced biofilm formation, while complementation restored wild-type phenotypes (Fig 4C), indicating RpoN2 is essential for flagellum-dependent motility and contributes to biofilm independently of DSF-QS system activation. RNA-seq at peak DSF-QS activity (14 h) identified 781 DEGs (423 downregulated, 358 upregulated, Fig 4D), with 276 (∼60%) overlapping the Clp regulon at the same time point (Fig 4E and S4). Most shared DEGs showed the same regulation, including motility and chemotaxis genes (down) and type IV pilins PilA/PilE (up), suggesting Clp control of motility and adhesion is largely mediated through RpoN2. Some genes, such as the chemotaxis protein CheW (Smlt2256) and flagellar proteins FliN (Smlt2281) and FlgH (Smlt2311), were uniquely downregulated in Δ*rpoN2* (Fig 4D). These findings provide a molecular explanation for the loss of motility observed in the K279a Δ*rpoN2* mutant and support the hypothesis that RpoN2 is at the top of the flagellar regulatory cascade in *S. maltophilia*. Regarding the loss of biofilm-forming ability of this mutant, although type IV pilins are overexpressed, the operon *smlt0706-0709*, which controls SMF-1 fimbriae production, is downregulated.

### The σ⁵⁴ paralog RpoN1 acts as a global regulator of fatty acid metabolism impacting DSF production in *S. maltophilia*

The K279a genome encodes two divergent σ⁵⁴ factor paralogs, Smlt1112 and Smlt2297, sharing 36% identity and retaining conserved σ⁵⁴ domains (S5 Fig). Smlt1112, which shares 73.7% identity with its *Xcc* ortholog RpoN1, is downregulated at the onset of stationary phase, whereas Smlt2297 (RpoN2) connects the DSF/Rpf-Clp axis to motility and adhesion. In *Xcc*, the σ⁵⁴ factor RpoN2 regulates flagellar biosynthesis, swimming motility, biofilm formation, EPS production, and virulence, whereas RpoN1 has a distinct role in FA metabolism and QS regulation [35]. Phylogenetic analysis separated RpoN proteins into two distinct groups within Pseudomonadota (Fig 5A): Group I, containing RpoN1 and canonical RpoN proteins from *E. coli* and *P. aeruginosa*, and Group II, containing RpoN2 orthologs. Similarly to *rpoN2*, *rpoN1* in the K279a genome is part of an operon encoding a σ⁵⁴-dependent activator (*smlt1111*) and a putative nitrogen regulatory protein (*smlt1110*). All *rpoN1* operon genes were also significantly downregulated in the wild-type strain at stationary phase entry (14 h) (S6 Fig and S1 File).

**Fig 5.**
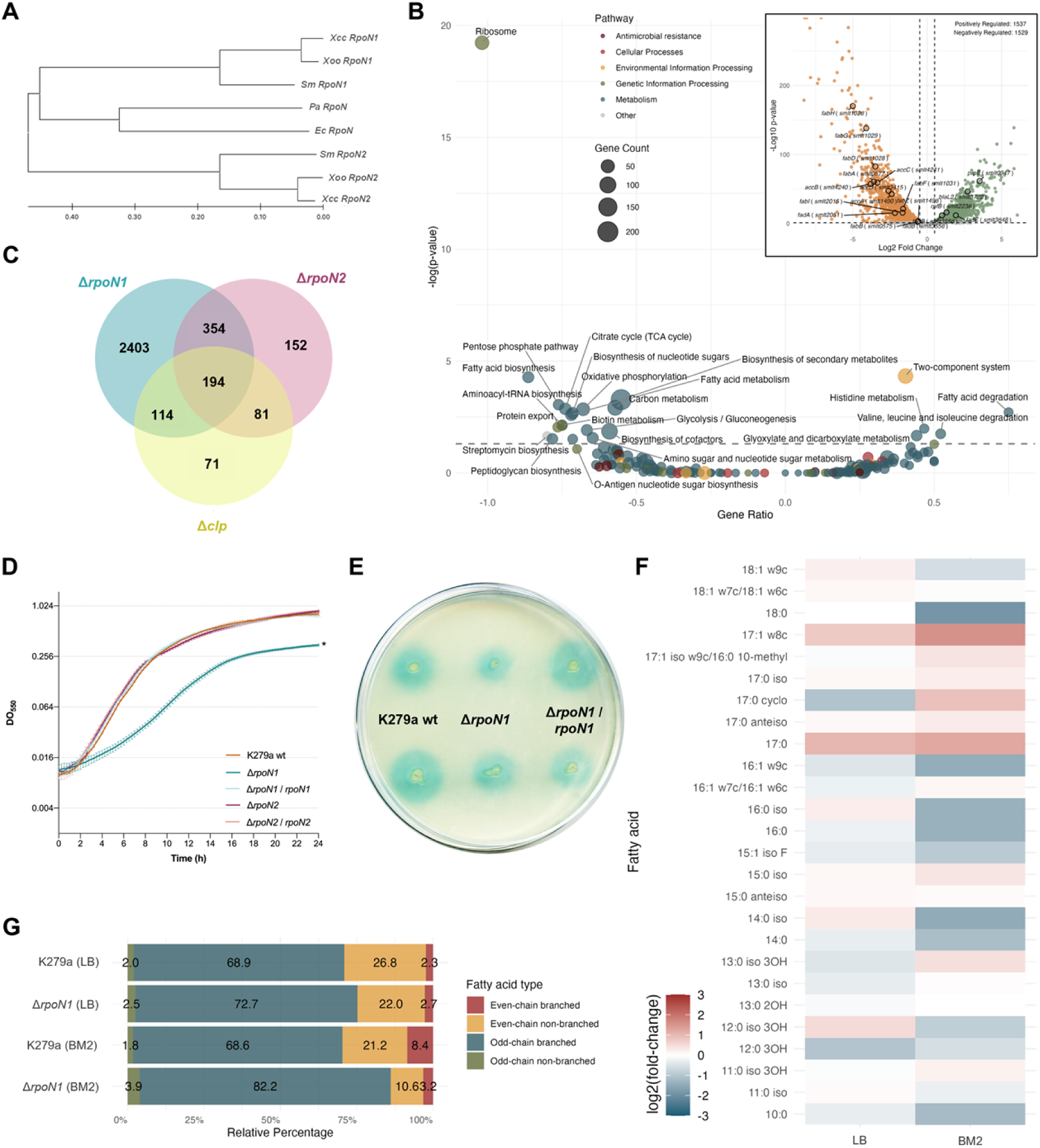
The σ⁵⁴ paralog RpoN1 acts as a global regulator of fatty acid metabolism and DSF production in *S. maltophilia* K279a. (A) Phylogenetic analysis of RpoN proteins showing two distinct clades: Group I (RpoN1 and canonical RpoN proteins) and Group II (RpoN2 proteins). The Neighbor-Joining tree was generated using MEGA-12 based on multiple sequence alignment. Abbreviations: *Xcc*, Xanthomonas campestris pv. *campestris*; *Xoo*, *Xanthomonas oryzae* pv. *oryzae*; *Sm*, *Stenotrophomonas maltophilia* K279a; Ec, *Escherichia coli*; Pa, *Pseudomonas aeruginosa*. The scale bar indicates the number of amino-acid substitutions per site. (B) KEGG pathway enrichment analysis of differentially expressed genes (DEGs) in the Δ*rpoN1* mutant at 14 h (DSF quorum-sensing activation). The x-axis shows gene ratio (negative, downregulated; positive, upregulated) and the y-axis shows pathway significance (−log₁₀ p). Circle size represents gene count and colours indicate pathway categories. The inset volcano plot displays DEGs (|log₂ fold change| ≥ 0.5, *p* < 0.05) with upregulated genes in green and downregulated genes in orange. Selected DEGs, including *blaL2* (*smlt3722*), *prpE* (*smlt0947*), and other genes related to fatty acid metabolism, are labelled and shown only when deregulated. (C) Venn diagram depicting overlap of DEGs among Δ*rpoN1*, Δ*rpoN2*, and Δ*clp* mutants at 14 h, highlighting extensive regulatory convergence. (D) Growth curves of wild-type K279a, Δ*rpoN1*, Δ*rpoN2*, and complemented strains in BM2 medium at 37 °C with agitation. Asterisks denote growth curves significantly different from wild type (*, *p* < 0.001; two-way ANOVA with Tukey’s test). (E) DSF production measured on NYG agar plates after 24 h at 28 °C using the *Xcc* DSF biosensor strain 8523/pL6engGUS. Blue coloration indicates DSF activity. (F) Heat map of cellular fatty acid (FA) composition in Δ*rpoN1* relative to wild type grown in LB or BM2 media. Only FAs comprising ≥0.2 % of total cellular FA content are shown; blue indicates reduction and red increase compared to wild type. (G) Stacked bar chart showing relative proportions of even-chain branched, even-chain non-branched, odd-chain branched, and odd-chain non-branched FAs in wild-type and Δ*rpoN1* cells grown in LB or BM2 media.

To assess functional specialization in *S. maltophilia*, particularly the role of RpoN1 in DSF-QS regulation, a K279a Δ*rpoN1* mutant was constructed for phenotypic analysis. RNA-seq of the K279a Δ*rpoN1* mutant at 14 h identified 3,065 DEGs (1,536 upregulated, 1,529 downregulated), representing 68% of the transcriptome (Fig 5B). KEGG analysis showed strong downregulation of core metabolic pathways, particularly the ribosome pathway, along with FA biosynthesis, pentose phosphate pathway, TCA cycle, protein export, oxidative phosphorylation, peptidoglycan biosynthesis, and amino sugar metabolism. In contrast, FA degradation, branched-chain amino acid degradation, glyoxylate and dicarboxylate metabolism, and two-component systems were upregulated, likely as compensatory responses (Fig 5B). The genes of the *rpf* cluster were also shown to be upregulated in mutant Δ*rpoN1*, highlighting the response regulator *rpfG* (+5.43-fold) and *rpfF* (+2.38-fold) and *rpfB* (+2.46-fold). This finding establishes for the first time a connection between the sigma factor RpoN1 and the *rpf* cluster.

The Δ*rpoN1* mutant shared ∼70% and 67% of its DEGs with the Δ*rpoN2* and Δ*clp* mutants, respectively (Fig 5C and S4), indicating substantial regulatory overlap and possible convergence among RpoN1, RpoN2, and Clp within the DSF-QS system. However, RpoN1 regulates numerous shared targets in the opposite direction, and uniquely controlled central metabolic pathways, suggesting a key role in metabolic adaptation and survival under nutrient limitation such as high cell density. Consistent with its metabolic role, the Δ*rpoN1* mutant showed a pronounced growth delay in the FA-limited BM2 medium (Fig 5D), unlike the Δ*rpoN2* mutant. It also produced significantly less DSF than the wild-type (Fig 5E). Additionally, ΔrpoN1 displayed a >16-fold increase in colistin sensitivity and an 8-fold increase in meropenem resistance (S2 Table), likely due to altered membrane composition and upregulation of BlaL2 (*smlt3722*, +6.56-fold). Complementation with *rpoN1* gene restored DSF production and antibiotic susceptibility to wild-type levels (Fig 5E and S2 Table).

To evaluate the role of RpoN1 in DSF production and lipid metabolism, FA profiles of wild-type and Δ*rpoN1* strains grown in LB and BM2 media were analysed (Fig 5F and S3 Table). In both strains, iso-15:0 and anteiso-15:0 were the major FAs, as reported previously [28,38]. Growth in BM2 increased even-chain BCFAs (branched-chain FAs) more than three-fold in the wild type, with a concomitant decrease in even-chain non-branched FAs (Fig 5G). In contrast, the Δ*rpoN1* mutant grown in BM2 exhibited a strong reduction in even-chain FAs, particularly BCFAs (>2-fold), while no differences were observed in LB (Fig 5G and S6 Table). These data indicate a key role for RpoN1 in even-chain FA synthesis, particularly BCFAs. Transcriptomic analysis (Fig 5B and S4) suggests that reduced acetyl-CoA availability in the mutant limits even-chain FA initiation, whereas upregulation of propionate-CoA ligase PrpE (*smlt0947*; +11.48-fold) may promote odd-chain FA synthesis [39], explaining the altered FA profile and reduced DSF production (see schematic in Fig 6). This shift likely affects membrane phospholipid composition and increases membrane vulnerability to membrane-disrupting agents. Collectively, our findings identify RpoN1 as a central regulator of lipid metabolism linked to quorum sensing and antibiotic resistance in *S. maltophilia*.

**Fig 6.**
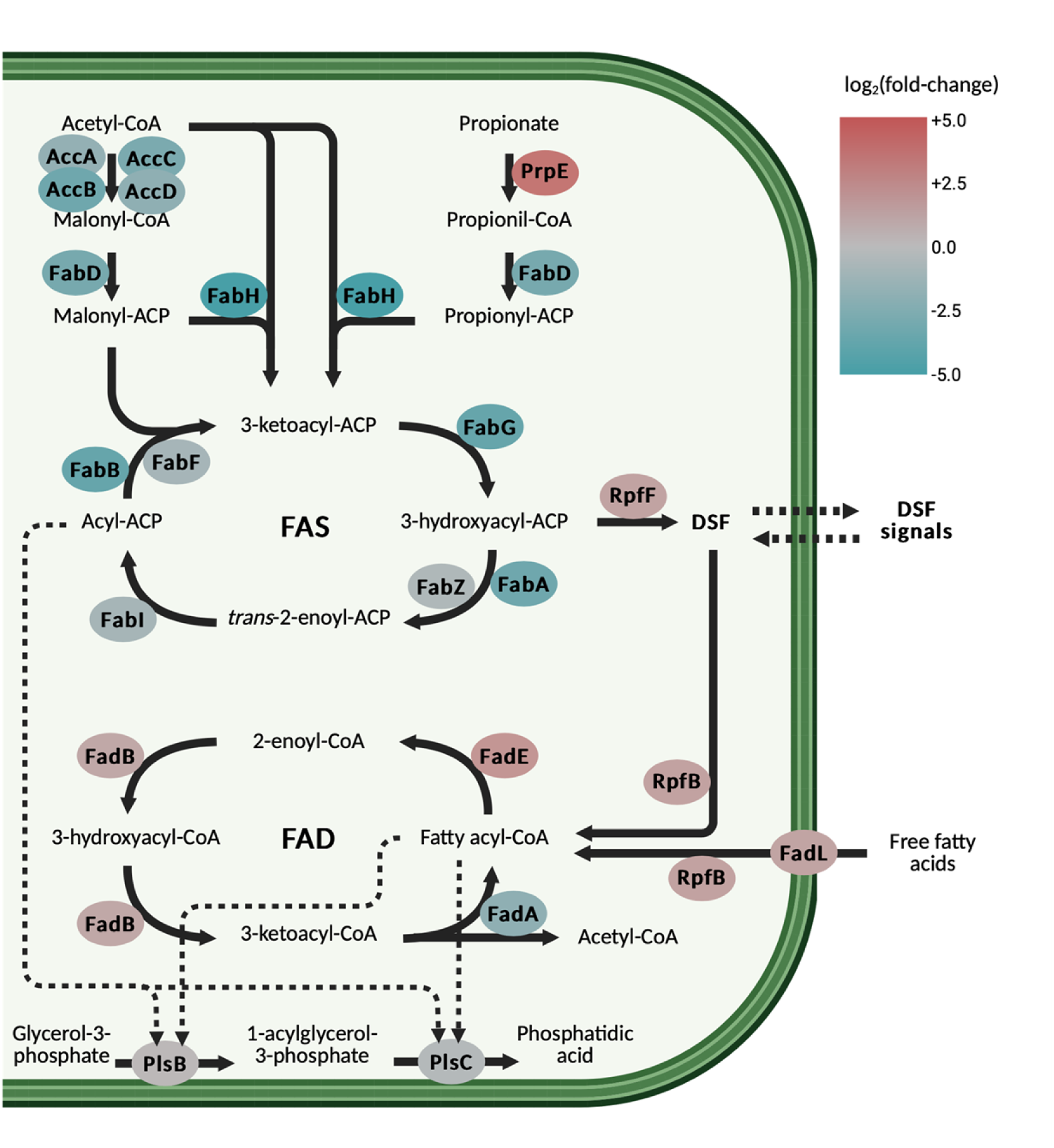
Integrated model of DSF/Rpf–Clp–RpoN regulatory interactions and metabolic control in *S. maltophilia* K279a. Schematic model illustrating how the σ⁵⁴ factor RpoN1 regulates fatty acid (FA) metabolism and DSF production. Both the fatty acid biosynthesis (FAS) and degradation (FAD) pathways, as well as the propionate pathway (PrpE), are shown. Enzymes are color-coded according to their log₂(fold-change) in the Δ*rpoN1* transcriptome relative to the wild type (blue, downregulated; red, upregulated). Downregulation of key FA biosynthetic enzymes together with upregulation of catabolic or propionate-related enzymes highlights a metabolic shift away from even-chain branched-chain FA synthesis toward FA degradation and odd-chain FA precursors derived from propionyl-CoA. Enzyme assignments are based on homology with functionally characterized proteins from related bacteria.

### RpfB coordinates global lipid metabolism beyond DSF turnover

Given its role in DSF homeostasis and its link to the RpoN1 regulon, we finally examined whether RpfB also contributes to global metabolic control. RpfB is annotated as a long-chain fatty acid-CoA ligase (FCL), an enzyme family that plays a crucial role in intermediary metabolism by activating free FAs into their CoA thioesters [14,40]. RpfB from *S. maltophilia* K279a shows high similarity to previously characterized FCL proteins, with 80% similarity to RpfB from *Xcc*, 71% to RpfB from *X. fastidiosa*, and 60% to FadD from *E. coli* (S7 Fig). A multiple sequence alignment revealed two highly conserved sequence elements corresponding to the ATP/AMP and FCL binding motifs, characteristic of adenylate-forming enzymes that share this catalytic function (S7 Fig). Although RpfB is known to modulate DSF turnover in other species [41,42], we aimed to explore its broader impact on bacterial physiology and metabolism reprogramming in *S. maltophilia*, extending beyond DSF-QS regulation.

The K279a Δ*rpfB* mutant was first grown in minimal medium supplemented with different carbon sources. The mutant exhibited normal growth in the presence of glucose but failed to grow when medium to long-chain FAs, including DSF, were used as the sole carbon source (Fig 7A). The wild-type and complemented strains can grow using FAs as the only source of carbon but with impaired fitness. We then analysed the FA composition of the K279a wild-type, the Δ*rpfB* mutant, and its complemented strain grown in rich medium (Fig 7B and S3 Table). The Δ*rpfB* mutant displayed a lower percentage of FAs with 16 or more carbon atoms (i.e., a higher percentage of FAs with fewer than 16 carbon atoms) relative to the total cellular FA content, compared to the wild-type strain (Fig 7C). The wild-type percentages were partially recovered in the complemented strain. These shifts likely result from the absence of RpfB acyl-CoA ligase activity, which would impair the reactivation of free long-chain FAs generated by the nonspecific thioesterase activity of RpfF and limit their reintegration into FA synthesis [14].

**Fig 7.**
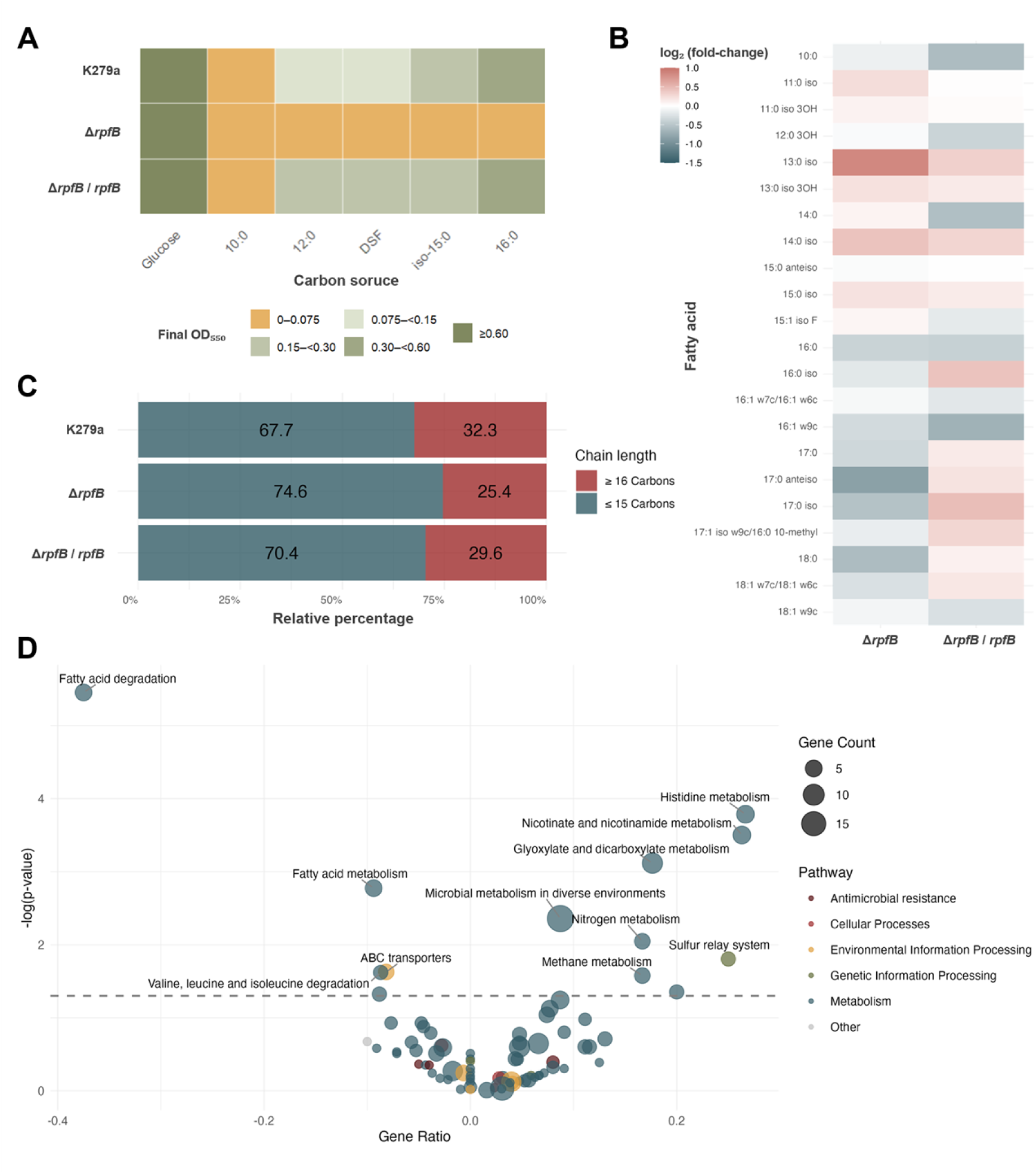
RpfB is required for fatty acid utilization and regulates global metabolism beyond DSF turnover in *S. maltophilia* K279a. (A) Categorical heat map summarizing growth of the K279a wild type, the ΔrpfB mutant, and the complemented strain in modified BM2 medium containing glucose or supplemented with individual fatty acids as the sole carbon source (decanoic acid [10:0], dodecanoic acid [12:0], cis-11-methyl-2-dodecenoic acid [DSF], 13-methyltetradecanoic acid [iso-15:0], or hexadecanoic acid [16:0]). Cultures were inoculated at OD₅₅₀ = 0.05 and incubated at 37 °C for 24 h with shaking. Tiles represent the mean OD₅₅₀ (n = 3) classified into discrete growth categories: no growth (OD₅₅₀ ≤ 0.075), low growth (0.075–<0.15), moderate growth (0.15–<0.30), high growth (0.30–<0.60), and very high growth (≥0.60). (B) Heat map of cellular fatty acid (FA) composition in Δ*rpfB* and complemented strains relative to wild type, showing log₂(fold-change) differences for FAs constituting >0.2 % of total content. Blue indicates reduction and red increase compared with wild type. (C) Distribution of cellular FAs by chain length in wild type, Δ*rpfB*, and complemented strains, illustrating depletion of long-chain (≥16 carbons) FAs in Δ*rpfB* and restoration in the complemented strain. (D) KEGG pathway enrichment analysis of differentially expressed genes (DEGs) in the Δ*rpfB* mutant at 14 h (DSF quorum-sensing activation). The x-axis shows gene ratio (negative, downregulated; positive, upregulated) and the y-axis shows pathway significance (−log₁₀ p). Circle size reflects DEG number and colours denote pathway categories. Significantly enriched pathways (*p* < 0.05, dotted line) are labelled.

The transcriptomic data for the Δ*rpfB* mutant (see Fig 2A) correlates with both an accumulation of DSF and an impaired ability of the mutant to utilize FAs as a carbon source. Functional enrichment analysis (GO/KEGG) revealed strong upregulation of energy production and conversion, transcription, and signal transduction terms, consistent with DSF overaccumulation and QS hyperactivation, while FA metabolism/degradation, branched-chain amino-acid catabolism, and transport-related processes were among the most significantly repressed categories (Fig 7D). In line with these global signatures, two β-oxidation loci, including *smlt2053-2051* and *smlt0264-0268*, were strongly downregulated. Collectively, these findings suggest that the absence of functional RpfB not only disrupts DSF turnover and quorum sensing regulation but also alters the bacterium’s overall metabolic state, triggering adaptive responses to meet energy demands and maintain cellular processes (see discussion).

## Discussion

Bacterial populations employ an array of sophisticated regulatory mechanisms to coordinate gene expression and adapt to diverse environmental pressures [43]. Among these, QS systems modulate processes central to survival, including biofilm formation, motility, virulence, and antibiotic resistance [44]. In *S. maltophilia* DSF-mediated QS shapes pathogen-host interactions, community-level behaviours, and environmental persistence [12]. Here, the DSF regulatory network drives reallocation of metabolic and physiological resources, particularly lipid flux, iron homeostasis, and envelope biogenesis, that underpins these behaviours and promotes adaptation. Beyond these broad roles, we reveal in *S. maltophilia* a previously unrecognized mechanism, absent from plant-pathogenic xanthomonads, in which DSF/Rpf-c-di-GMP signalling directly controls σ⁵⁴-dependent regulation, coupling quorum-sensing to lifestyle and metabolic reprogramming in a human opportunistic pathogen.

The observed coupling between entry into stationary phase, DSF system activation, and a decrease in intracellular c-di-GMP in *S. maltophilia* is consistent with the RpfC-RpfG architecture established in *Lysobacteraceae*, where DSF-bound RpfC activates the HD-GYP phosphodiesterase RpfG to lower c-di-GMP [18,45–47]. Downstream, Clp mediates this coupling at the transcriptional level. Motif discovery identified a CRP-like palindromic sequence (TGTGA-N₆-TCACA) enriched among Clp-activated targets, matching the consensus Clp site in *X. campestris* [16,33], and biochemical assays show that c-di-GMP directly inhibits Clp-DNA binding. Reporter strains driven by a Clp motif-containing promoter confirmed the inverse *in vivo* relationship between c-di-GMP and Clp-dependent transcription, supporting a DSF/Rpf-Clp axis that reprograms transcription across growth phases (Fig 8). Notably, the Clp regulon shifts with physiology: at exponential phase (rising c-di-GMP), many DEGs lack Clp motifs and appear indirectly controlled; at QS peak, motif-bearing targets predominate, and functions related to secretion, envelope, and surface structures are primarily activated, a regulatory configuration consistent with growth phase-dependent transcriptional reprogramming.

**Fig 8.**
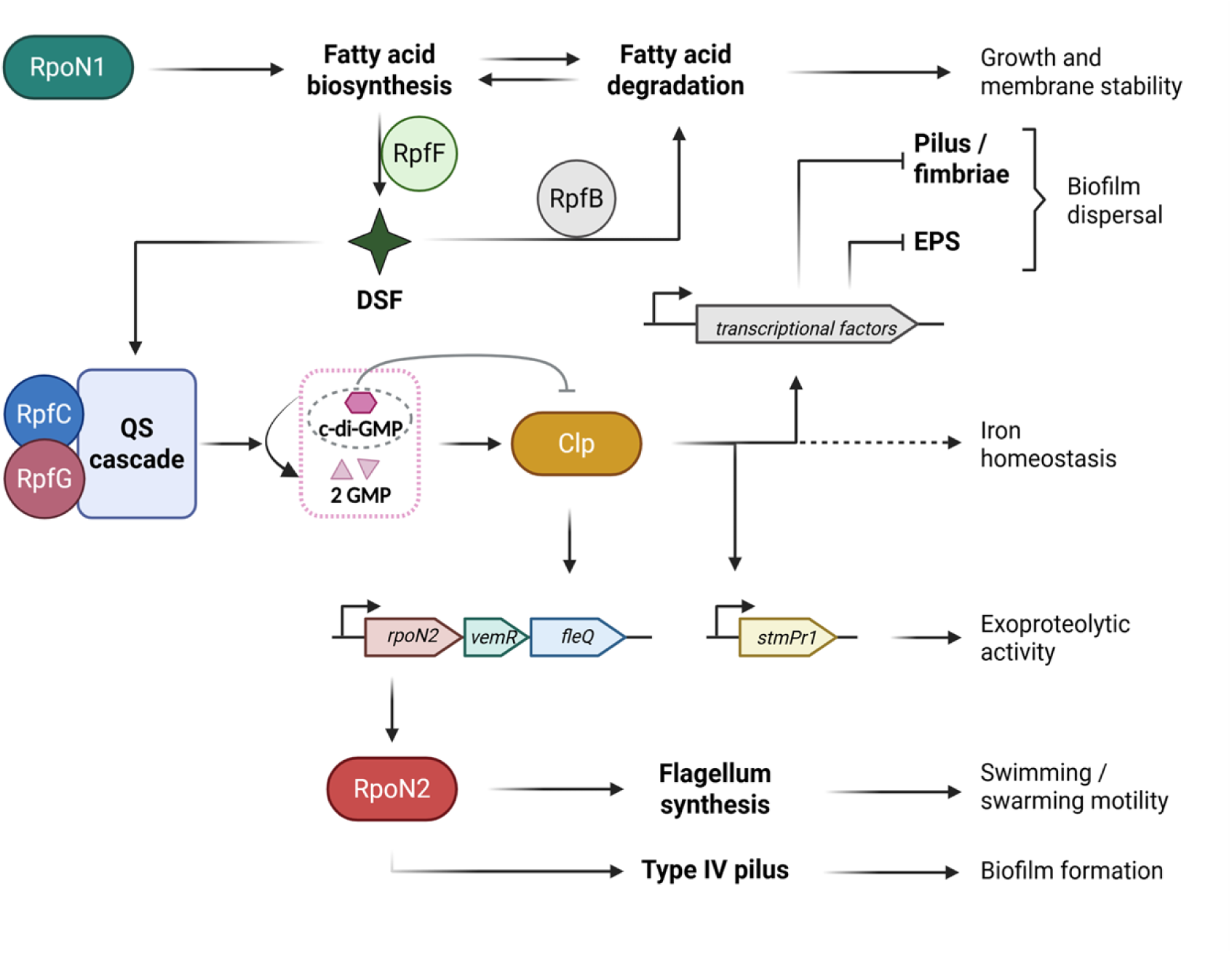
Integrated schematic representation summarizing the DSF/Rpf–Clp–RpoN regulatory network. The diagram depicts how DSF signaling, mediated by the Rpf system, regulates intracellular c-di-GMP levels and modulates Clp-dependent transcription. Clp in turn controls σ⁵⁴ (RpoN2)-dependent motility and virulence genes, while RpoN1 coordinates fatty acid metabolism and DSF biosynthesis, linking quorum sensing, metabolism, and the planktonic–biofilm lifestyle transition. Dashed lines indicate indirect or putative regulatory interactions supported by transcriptomic evidence.

The regulatory pathway DSF/Rpf-Clp extends to a σ⁵⁴-dependent circuit in *S. maltophilia*. Clp directly binds the promoter of the operon *rpoN2-smlt2296-fleQ*, and c-di-GMP disrupts this interaction, identifying Clp as a control point for RpoN2-dependent transcription that operates when c-di-GMP is low. Δ*rpoN2* abolishes swimming and swarming and reduces expression of flagellar and chemotaxis genes, consistent with σ⁵⁴-dependent regulation described in *X. campestris* [35,36], whereas *fleQ* and *fliA* transcript levels remain largely unchanged, like *P. aeruginosa*, where FleQ functions as a master enhancer binding proteins (EBP) with constitutive expression [48]. In *S. maltophilia*, however, *fleQ* is encoded in the same operon as *rpoN2* and is partially controlled by Clp, indicating a species-specific point of integration between QS, c-di-GMP signalling, and the flagellar transcriptional cascade (Fig 8). This organization supports a wiring logic distinct from plant-pathogenic xanthomonads, aligning σ⁵⁴ control more tightly with DSF status in *S. maltophilia* [49]. Functionally, a reciprocal increase in type IV pili/SMF-1 when flagellar gene expression is reduced is consistent with a DSF-mediated model that balances locomotion and adhesion under c-di-GMP control, with these phenotypes arising from metabolic reallocation. In other pathogenic bacteria c-di-GMP is considered a switch molecule that regulates the lifestyle transition from motility to biofilm [27].

Finally, the *smlt2296* operon gene shares 92% identity with the response regulator VemR of *Xcc*, which modulates ∼10% of the bacterial genome, including genes involved in exopolysaccharide and exoenzyme production, motility, and virulence [49,50].

QS also intersects iron homeostasis. At the peak of QS activity, Fur-regulated systems as stenobactin biosynthesis/export (*entCEBB’FA*), ferric-citrate and ferrous iron uptake (*fciTABC/feoABI*), and heme acquisition are induced, consistent with iron limitation at high cell density [51–53]. Prior studies show that iron limitation increases DSF production and that *fur* mutants overproduce DSF even under iron-replete conditions, indicating bidirectional coupling between DSF synthesis and Fur-mediated regulation [54]. The enrichment of Fur-box-like sequences among promoters upregulated in Δ*clp* at the QS peak suggests that Clp influences iron homeostasis indirectly, most likely by altering intracellular iron availability rather than by direct promoter binding. These observations indicate that DSF-dependent signalling coordinates iron uptake and utilization with cell-surface and secretion functions, a hallmark of metabolism reprogramming in mediating bacterial virulence.

RpfB is positioned at the interface between signal turnover and lipid homeostasis (Fig 6 and 8). Deletion of *rpfB* in *S. maltophilia* K279a not only impacts DSF turnover but also prevents growth on medium- and long-chain FAs as sole carbon sources, shifts the cellular fatty-acid distribution toward shorter chains (reduced ≥C16 species), and increases susceptibility to colistin. Transcriptome analysis of Δ*rpfB* shows broad upregulation of transcription and signal-transduction functions and downregulation of fatty-acid metabolism and β-oxidation, including repression of two β-oxidation loci (*smlt2053-2051* and *smlt0264-0268*) that resemble the TetR-FadB-FadA organization in *Xanthomonas citri* and PsrA-regulated modules in *P. aeruginosa* [55,56]. In *S. maltophilia*, the TetR-type regulator Smlt2053 has been proposed to act as an intracellular sensor for DSF, coordinating signal turnover by promoting flux of excess long-chain FAs into β-oxidation [28]. Mechanistically, these observations are consistent with a recycling model in which RpfB counteracts the nonspecific thioesterase activity of RpfF by reconverting liberated long-chain FAs [41], including DSF, into acyl-CoA, thereby maintaining the pool of long acyl chains required for outer-membrane lipid composition and limiting susceptibility to polymyxins. The opposite colistin phenotype of Δ*rpfF* further supports that the Δ*rpfB* susceptibility arises from lipidomic imbalance rather than from loss of QS activation. In *S. maltophilia*, these lipidic adjustments connect DSF turnover to envelope physiology and drug response, extending DSF biology into clinically relevant terrain less emphasized in plant-pathogen models [57].

Finally, a link between central-carbon metabolism, DSF output, and outer-membrane susceptibility in *S. maltophilia* has been highlighted with this work (Fig 6). First, a functional specialization between alternative σ⁵⁴ paralogs links motility regulation with fatty-acid precursor supply. Consistent with observations in *Xanthomonas spp.*, RpoN2 controls flagellar gene expression, whereas RpoN1 regulates central metabolic pathways [58,59]. In *S. maltophilia* Δ*rpoN1*, pathways for ribosome biogenesis, the TCA cycle, oxidative phosphorylation, peptidoglycan biosynthesis, and fatty-acid biosynthesis are downregulated, while fatty-acid and amino acids degradation are upregulated, a pattern consistent with resource-conservation responses described in *Xcc* grown under lipid limitation [35]. Lipidomic and transcriptomic data from the RpoN1-deficient strain indicate reduced acetyl-CoA availability alongside PrpE upregulation, favouring propionyl-CoA-initiated FA synthesis; this is consistent with the established role of propionyl-CoA in initiating odd-chain FAs [60]. Here this is associated with lower levels of even-chain branched FAs and reduced DSF production and is accompanied by membrane remodelling and increased susceptibility to colistin. Notably, Li *et al*. showed in *X. campestris* that RpoN1 influences branched-chain fatty-acid pools and modulates DSF-family output, but the direction and route differ from our observations: in *S. maltophilia*, RpoN1 constrains DSF through reduced acetyl-CoA availability and a shift toward propionyl-CoA-initiated chains (lower even-chain BCFAs), linking precursor economy to DSF synthesis and membrane remodelling, whereas in *X. campestris* Δ*rpoN1* increased DSF-family signals alongside altered BCFAs [35].

Importantly, *rpoN1* expression decreases at the onset of stationary phase in the wild-type strain, coinciding with the induction of *rpfF* and maximal DSF production, whereas deletion of *rpoN1* results in increased transcription of *rpfF*, *rpfG* and *rpfB* at 14 h. Given that *rpfF* is monocistronic and transcriptionally independent of *rpfB*, these observations suggest that RpoN1 does not act as a simple direct repressor of the *rpf* cluster but rather constrains DSF signalling primarily through metabolic control. Under this model, high RpoN1 activity during exponential growth limits DSF synthesis by restricting fatty-acid precursor availability, whereas downregulation of *rpoN1* upon entry into stationary phase relieves this constraint, allowing *rpfF* induction and DSF production. However, increased *rpfF* transcription in the Δ*rpoN1* background does not translate into elevated DSF levels, likely reflecting an imbalance between DSF synthesis and the availability of appropriate lipid substrates and signal turnover efficiency (*rpfB* increases its expression as well).

The Δ*rpfB* mutant (DSF overproducer) also exhibited increased colistin sensitivity, whereas the Δ*rpfF* mutant showed the opposite phenotype, suggesting altered membrane phospholipids in these strains and a link to DSF levels as we previously proposed [10]. Changes in the phospholipid fatty-acyl ratio are known to modulate susceptibility to membrane-damaging agents such as polymyxins [61,62]. In *Pseudomonas spp.*, a single copy of the σ⁵⁴ factor RpoN regulates the flagellin gene as well as several metabolic processes, particularly those involving amino acid and nitrogen metabolism and QS-related pathways [63–65]. The evolutionary duplication of the *rpoN* gene into two σ⁵⁴ factors with distinct functions has been maintained in *S. maltophilia*, most likely to adapt the bacteria to distinct niches in the colonization and pathogenesis process. Together with the Clp-dependent control of *rpoN2-vemR-fleQ*, these results support a species-specific integration of DSF signalling with σ⁵⁴ specialization in *S. maltophilia*.

Taken together, these findings indicate that DSF-mediated cell-cell signalling in *S. maltophilia* consistently drives metabolic reprioritization. The DSF signalling cascade integrates cell-density cues with second messenger c-di-GMP dynamics and σ⁵⁴ (RpoN2) transcriptional control through the global regulator Clp, coupling quorum state to motility, while an RpoN1-centred metabolic arm supplies fatty-acid precursors that sustain DSF biosynthesis and RpfB links signal turnover to acyl-CoA pools (Fig 8). This circuitry connects behaviour to metabolism, clarifies differences with *Xanthomonas* models and frames clinically relevant phenotypes in *S. maltophilia*. DSF homeostasis metabolic processes also contribute to shaping pathogenic phenotypes such as biofilm formation and antibiotic resistance. On one hand σ⁵⁴ factor RpoN1 governs lipid and energy precursor supply that supports membrane architecture and DSF synthesis while RpfB closes the loop by coupling signal turnover to fatty-acyl-CoA activation. This network architecture exemplifies metabolism reprogramming strategies, in which QS signals, second messengers, and core metabolic pathways are coordinated to promote persistence, dissemination, and stress tolerance in host-relevant conditions. However, the regulatory network appears considerably more complex, as transcriptomic analyses of individual mutants reveal numerous additional genes and mechanisms that may be governed by Clp-independent signalling pathways. These insights underscore the need for deeper investigation, including the influence of iron through Fur on quorum-sensing outputs and the broader regulatory architecture.

## Materials and methods

### Bacterial strains and growth conditions

All bacterial strains used in this study are listed in S4 Table. *S. maltophilia* K279a was employed as the wild-type strain and as a representative of the *rpf*-1 group [66]. *Escherichia coli* DH5α, SY327, and SM10 (λPir) were used for cloning purposes, with SY327 and SM10 specifically employed for plasmids carrying an R6Kγ origin of replication. *E. coli* BL21(DE3) was used for heterologous protein expression. The DSF biosensor strain was *Xanthomonas campestris* pv. *campestris* (*Xcc*) 8523, a Δ*rpfF* mutant carrying a chromosomal fusion of the *engXCA* promoter to *gusA* on plasmid pLAFR6 (pL6engGUS), which enables DSF detection [18].

Unless otherwise indicated, *S. maltophilia* and *E. coli* strains were routinely grown in Miller’s LB medium (10 g/L tryptone, 5 g/L yeast extract, 10 g/L NaCl) at 37 °C with shaking at 250 rpm, or on LB supplemented with 1.5% (w/v) agar. *E. coli* SM10 (λPir)/pUX-BF13 was cultured at 30 °C, and *Xcc* 8523 was maintained at 28 °C in NYG medium (0.5% peptone, 0.3% yeast extract, 2% glycerol) supplemented with tetracycline (10 µg/mL). For phenotypic assays, modified BM2 minimal medium was used [67], consisting of 62 mM potassium phosphate buffer (pH 7.2), 2 mM MgSO₄, 10 µM FeSO₄, 0.4% (w/v) D-glucose, and 0.5% (w/v) casamino acids. Growth was routinely monitored by measuring the optical density at 550 nm (OD₅₅₀) using a Novaspec II spectrophotometer (Pharmacia) or a Multiskan FC microplate reader (Thermo Fisher Scientific).

### Construction of markerless deletion mutants and genetic complementation

Markerless deletion mutants were generated using the pGPI-SceI / pDAI-SceI-SacB two-step allelic exchange system [68], which was previously adapted for *S. maltophilia* [28,66]. Approximately 700 bp of DNA upstream and downstream of each target gene were PCR-amplified and cloned into the non-replicative pGPI-SceI-XCm vector (see plasmids and primers on S5 and S6 Tables), which carries a chloramphenicol resistance gene, the *xilE* reporter, and a unique I-SceI recognition site. Cloning was performed by NEBuilder® HiFi DNA Assembly, and constructs were verified by colony PCR and Sanger sequencing in *E. coli* SY327.

The resulting suicide plasmids were transferred to *S. maltophilia* K279a via triparental mating using *E. coli* DH5α/pRK2013 as the helper strain. Transconjugants in which the plasmid integrated by homologous recombination were selected on LB agar containing chloramphenicol (60 µg/mL) and norfloxacin (5 µg/mL). Integration was further confirmed by spraying colonies with 0.45 M pyrocatechol, which produces a yellow colour in the presence of *xilE*. To resolve the cointegration, plasmid pDAI-SceI-SacB expressing the I-SceI endonuclease was introduced by electroporation (2.5 kV, 200 Ω, 25 µF) [69]. This induced a double-strand break, stimulating a second recombination event that resulted either in excision of the plasmid with restoration of the wild-type allele or in a markerless deletion of the target gene. Recombinants were selected on LB agar with tetracycline (60 µg/mL), and deletion events were verified by PCR using external primers and sequencing. The pDAI-SceI-SacB plasmid was cured by counter-selection on LB agar containing 5% sucrose, exploiting the lethality conferred by the *sacB* gene, and loss of tetracycline resistance confirmed plasmid removal.

For genetic complementation, the deleted gene together with its own promoter and regulatory sequences was cloned into the broad-host-range vector pBBR1-MCS1. When tight control of expression or operon structure required an alternative strategy, the arabinose-inducible derivative pBBR1-pBAD-MCS1 was used [70]. Recombinant plasmids were verified by PCR and sequencing, introduced into *S. maltophilia* mutants by electroporation, and selected on LB agar supplemented with chloramphenicol (60 µg/mL).

### Fluorescent c-di-GMP biosensor construction and in vivo sensing

A mini-Tn7 delivery system optimized for *Lysobacteraceae* [71,72] was used to integrate a fluorescent reporter at the neutral chromosomal site downstream of the *glmS* gene. The reporter plasmid pUC18T-mini-Tn7T-Gm-PrpoD-sfGFP, which drives sfGFP expression from the *rpoD* promoter, was modified by replacing the promoter with the *smlt3461* promoter (PCR-amplified and inserted at KpnI/BamHI sites), yielding pUC18T-mini-Tn7T-Gm-P*smlt3461*-sfGFP.

Four-parental matings were carried out with *E. coli* DH5α/pUC18T-mini-Tn7T-Gm-P*smlt3461*-sfGFP (donor), DH5α/pRK2013 (helper), SM10/pUX-BF13 (helper), and *S. maltophilia* recipient cells. Transconjugants were selected on LB agar containing gentamicin (60 µg/mL) and norfloxacin (5 µg/mL), and correct chromosomal integration was confirmed by PCR using primers PTn7L/Smlt4098-Ctrl and PTn7R/Smlt4099-Ctrl (S6 Table).

For in vivo c-di-GMP quantification, cultures of the reporter strain were sampled at different growth phases, and sfGFP fluorescence was measured using a Victor V3 1420 multilabel plate reader (PerkinElmer) with excitation at 485 nm and emission at 535 nm. Reporter fluorescence inversely correlated with intracellular c-di-GMP concentrations.

### Recombinant Smlt4306 (Clp) production and purification

The coding sequence of *clp* (*smlt4306*) was amplified from *S. maltophilia* K279a and cloned into pET28b to produce N-terminally His₆-tagged Clp (pET28b-Clp-H6). The plasmid was introduced into *E. coli* BL21(DE3), which was cultured in 6 L of LB medium with kanamycin (50 µg/mL) at 37 °C until OD₅₅₀ reached 0.5. Protein expression was induced with 1 mM IPTG and continued for 3 h. Cells were harvested, washed with PBS, and lysed by two passages through an Emulsiflex C5 homogenizer followed by mild sonication in binding buffer (20 mM Tris-HCl pH 8.0, 250 mM NaCl, 100 mM LiCl, 1% glycerol, 1 mM DTT, 20 mM imidazole). The clarified lysate was filtered (0.22 µm) and subjected to nickel-affinity chromatography on a HisTrap HP column (Cytiva) using a linear imidazole gradient up to 500 mM. Eluted protein was dialyzed against 20 mM Tris-HCl pH 8.0, 250 mM NaCl, 100 mM LiCl, 1% glycerol, and 1 mM DTT. Purity and identity were confirmed by SDS-PAGE, western blotting, and mass spectrometry. Protein purification was done by the Protein Production Platform (U1) Unit of CIBER in Bioengineering, Biomaterials & Nanomedicine CIBER-BBN, Spain (https://www.nanbiosis.es/platform-units/).

### Quantification of c-di-GMP by ELISA

Intracellular c-di-GMP levels were determined using the Cayman c-di-GMP ELISA kit. Cells were harvested at defined growth phases, washed three times with cold PBS, and 8 × 10⁹ cells were lysed in 500 µL of B-PER reagent (Thermo Fisher) for 15 min at room temperature. Lysates were clarified by centrifugation (15,000×g, 10 min, 4 °C), and protein concentration was measured by the Pierce™ BCA Protein Assay (Thermo Scientific). c-di-GMP concentrations were quantified following the manufacturer’s instructions and normalized to total protein.

### Cellular fatty acid analysis

Cultures grown for 24 h at 37 °C in LB or BM2 were analyzed at the Spanish Type Culture Collection (CECT, University of Valencia) using the MIDI Microbial Identification System and an Agilent 6850 gas chromatograph with the Sherlock TSBA6 method [28].

### DSF detection

DSF production was measured using *Xcc* 8523 pL6engGUS as reporter [9]. Reporter cultures (OD₅₅₀ = 0.7) were mixed to a final OD₅₅₀ of 0.07 with NYG containing 1% noble agar and X-Glu (80 µg/mL), poured into plates, and inoculated with test strains. After 24 h at 28 °C, blue halos indicated DSF activity.

### Electrophoretic mobility shift assay (EMSA)

Promoter fragments (150-300 bp) were incubated with purified Clp in buffer (10-40 mM Tris-Glycine, 20 mM KCl, 7.5% glycerol, 0.01 mg/mL BSA, 1 mM DTT, 20 ng/µL poly(dI·dC)) with or without c-di-GMP [73]. After 30 min at room temperature, mixtures were resolved on 6% polyacrylamide gels in 0.5× TBE at 4 °C, stained with RedSafe™, and visualized with a VersaDoc™ imaging system (Bio-Rad).

### Phenotypic assays

#### Growth curves

Overnight cultures were diluted to an OD₅₅₀ of 0.01 and inoculated in triplicate into 96-well plates containing LB or BM2. Plates were incubated at 30 °C or 37 °C in a Multiskan FC reader with continuous shaking, and OD₅₅₀ was recorded every 15 min for 24 h.

#### Biofilm formation

Biofilm production was quantified by crystal violet staining [74]. Overnight cultures in BM2 were adjusted to an OD₅₅₀ of 0.05 and inoculated into 96-well plates (200 µL/well), incubated statically at 30 °C or 37 °C for 24 h, washed, heat-fixed (60 °C, 15 min), and stained with 0.1% crystal violet for 15 min. After washing and drying, the bound dye was solubilized with 30% acetic acid and quantified at OD₅₅₀.

#### Exopolysaccharide production

Colonies were grown either on BHI agar supplemented with 3% sucrose and Congo red (8 mg/mL) at 37 °C for 24 h, or on salt-free LB agar (1% agar) supplemented with Congo red (40 µg/mL) and Coomassie Brilliant Blue (20 µg/mL) at 30 °C for several days [75,76]. Colony morphology and dye binding were assessed visually. Exopolysaccharide (EPS) producing strains typically produce red colonies with dark red spots due to absorption of the Congo red dye [72].

#### Exoproteolytic activity

LB agar supplemented with 2% skim milk or 5% egg yolk was inoculated and incubated for 24 h at 37 °C; protease activity was inferred from clear halos around colonies.

#### Motility

Swimming motility was tested on LB with 0.25% noble agar and swarming on BM2 with 0.5% noble agar as described [77]. Inoculated plates were incubated at 30 °C for 48 h (swimming) or 5 days (swarming), and migration diameters were measured.

#### Antimicrobial susceptibility

Minimum inhibitory concentrations (MICs) were determined by broth microdilution according to CLSI guidelines and EUCAST recommendations for colistin [78,79]. Cation-adjusted Mueller-Hinton broth (25 µg/mL Ca²⁺ and 12.5 µg/mL Mg²⁺) was used. Two-fold antibiotic dilutions were prepared in 96-well plates and inoculated with 5 × 10⁵ CFU/mL. After 24 h at 37 °C, MICs were recorded visually and confirmed with resazurin (0.01% w/v).

#### Scanning electron microscopy

Cells from overnight LB cultures were fixed in 1% paraformaldehyde, dehydrated through an ethanol series, critical-point dried (Balzers CPD 030), sputter-coated with gold (SCD 050), and imaged using a Zeiss Merlin SEM.

### RNA-seq transcriptomics

Triplicate cultures of *S. maltophilia* K279a were inoculated at OD₅₅₀ = 0.05 in LB and grown at 37 °C with shaking. Cells were harvested after 5 or 14 h depending on the experimental design. Total RNA was extracted with the RNeasy Mini Kit (QIAGEN) and treated on-column with DNase I. RNA quality was verified using an Agilent 2100 Bioanalyzer, and only samples with RIN > 8 were used for library construction and Illumina sequencing at Novogene (Beijing, China).

Adapter trimming and quality filtering of FASTQ reads were performed using fastp. Clean reads were aligned to the *S. maltophilia* reference genome (NCBI) with Bowtie2 [80]. Gene-level counts were obtained with FeatureCounts, and FPKM values were calculated. Differential expression between conditions (three biological replicates each) was determined using DESeq2 [81], applying the Benjamini-Hochberg procedure to control false discovery. Genes with adjusted *p* (*p*) < 0.05 and |log₂ fold change| > 0.5 were considered differentially expressed. Gene Ontology (GO) [82,83] and KEGG pathway [84,85] enrichment analyses were performed with the clusterProfiler R package [86]. Data visualization used ggplot2 (https://link.springer.com/book/10.1007/978-3-319-24277-4), circlize [87], UpSetR [88], and VennDiagram [89].

### Bioinformatics analysis

Genomic sequences were retrieved from the NCBI RefSeq database and analyzed with BLAST. The Artemis genome browser was used to visualize genomic features and mapped RNA-seq reads [90]. Multiple sequence alignments were performed with ClustalW in MEGA12 [91]. Motif analysis was conducted using the MEME Suite, TomTom was used to identify known motifs, and FIMO to locate specific motif occurrences [92]. Regulatory networks were explored using RegPrecise 3.0 [93].

### Statistical analysis

Normality of data was assessed by the Shapiro-Wilk test. For normally distributed data, one-way ANOVA was used to compare three or more groups differing by a single factor, while two-way ANOVA was applied when assessing two independent variables, such as treatment and time in growth curves. Statistical significance was defined at *p* < 0.05.

### Data Availability

The RNA-seq data generated in this study have been deposited at the GEO SuperSerie GSE294404 (SubSeries GSE294324 and GSE294403). Source data are provided with this paper.

## Supporting information captions

**S1 Figure.**
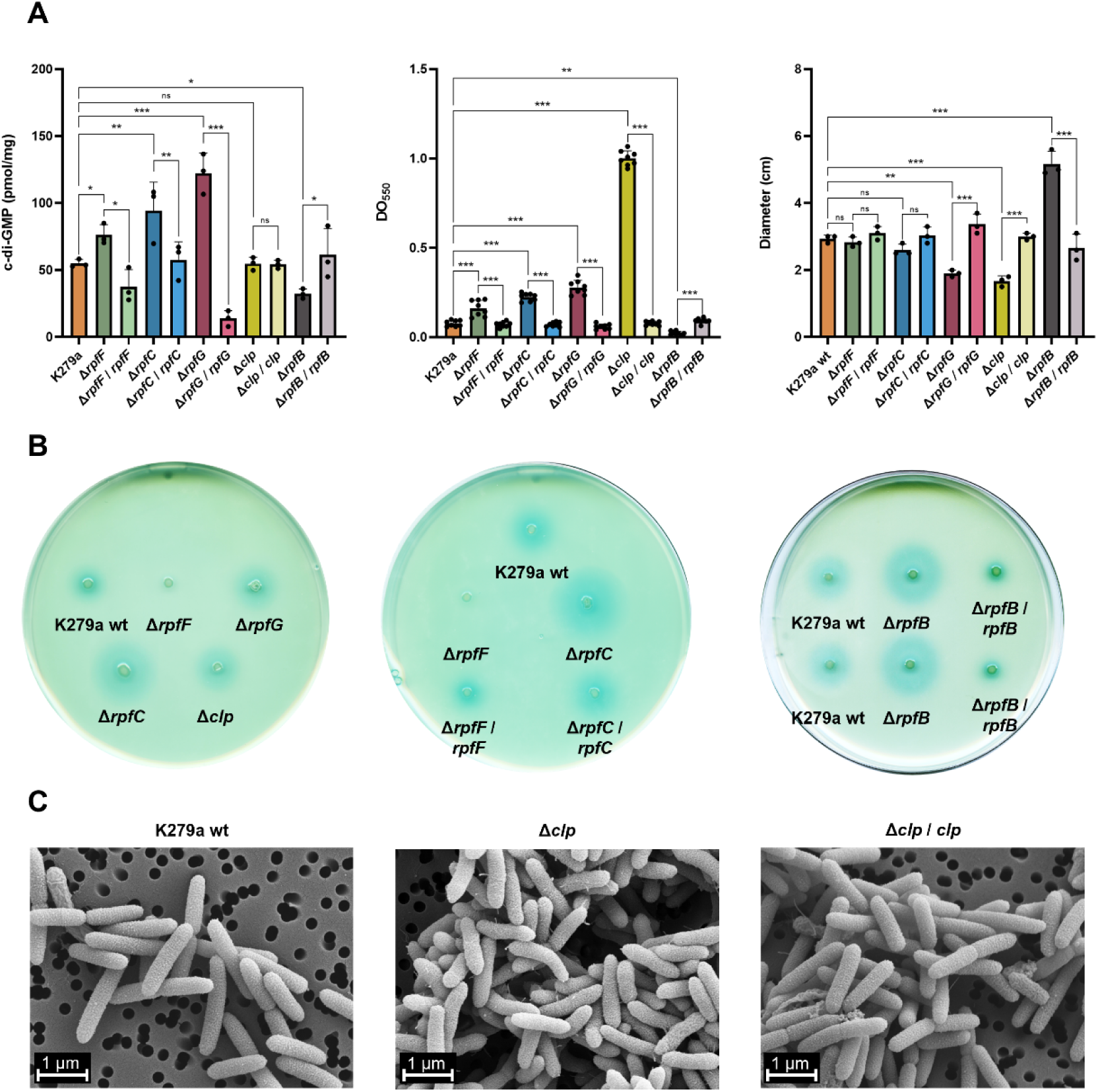
Functional analysis of the DSF/Rpf–Clp signaling cascade in *S. maltophilia* K279a. (A) Quantification of intracellular c-di-GMP levels in *S. maltophilia* K279a wild type, deletion mutants (Δ*rpfF*, Δ*rpfC*, Δ*rpfG*, Δ*rpfB*, and Δ*clp*), and their respective complemented strains after 14 h of growth in LB at 37°C using the Cayman’s cyclic-di-GMP ELISA kit. Biofilm formation was assessed in BM2 glucose minimal medium supplemented with casamino acids at 37°C for 24 h. Swimming motility was measured on 0.25% LB agar at 30°C after 48 h. Bars represent mean values ± SD from at least three biological replicates. Statistical significance was evaluated using one-way ANOVA with Tukey’s post hoc test (*, *p* < 0.05; **, *p* < 0.01; ***, *p* < 0.001; *ns*, not significant). (B) DSF production was visualized by a colony bioassay on NYG agar at 28°C for 24 h, using the biosensor strain X. campestris pv. campestris 8523/pL6engGUS; blue coloration indicates DSF activity. (C) Scanning electron micrographs of *S. maltophilia* K279a wild-type, Δ*clp* mutant, and complemented strain grown in LB liquid medium until 14 h (peak DSF-QS activity). The Δ*clp* mutant displays abundant pili and fimbriae structures, absent in wild-type and complemented strains. Scale bar, 1 μm.

**S2 Figure.**
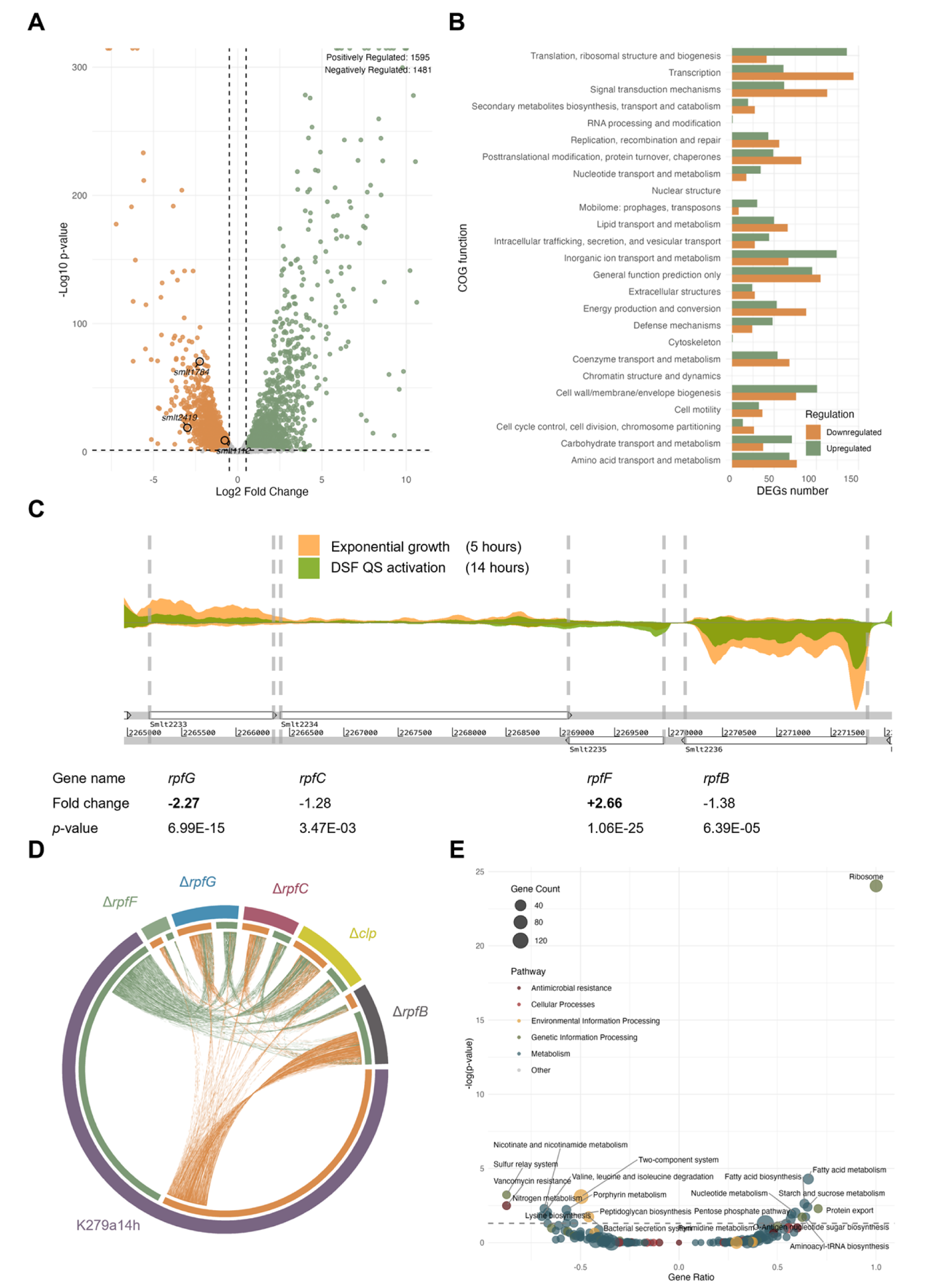
Growth phase–dependent transcriptomic changes in wild-type *S. maltophilia* K279a. (A) Volcano plot showing differentially expressed genes (DEGs) in wild-type K279a at 14 h (stationary phase/DSF-QS activation) relative to 5 h (exponential growth). Genes with |log₂ fold change| ≥ 0.5 and p < 0.05 are shown (green, upregulated; orange, downregulated). (B) Functional classification of DEGs into clusters of orthologous groups (COG). The x-axis shows the number of genes in each COG category and the y-axis the functional groups. Upregulated and downregulated genes are indicated using the same colour scheme as in panel A. Genes of unknown function were excluded. (C) RNA-seq read coverage across the *rpf* gene cluster, comparing samples at 14 h with those at 5 h. Fold-change values indicate transcriptional variation between the two conditions; significant changes are highlighted in bold. (D) Circos plot showing DEGs shared between wild-type K279a at 14 h and Δ*rpfF*, Δ*rpfC*, ΔrpfG, Δclp, and ΔrpfB mutants. The plot highlights transcriptional convergence and divergence within the DSF/Rpf–Clp axis and growth phase regulation in the wild type. Green lines denote downregulated genes and orange lines upregulated genes. (E) KEGG pathway enrichment analysis of DEGs in wild-type K279a at 14 h relative to 5 h. The x-axis shows the gene ratio (negative values, downregulated pathways; positive values, upregulated pathways) and the y-axis shows pathway significance (−log₁₀ p). Circle size reflects the number of DEGs, and colours indicate pathway categories. Significantly enriched pathways (p < 0.05, dotted line) are labelled.

**S3 Figure.**
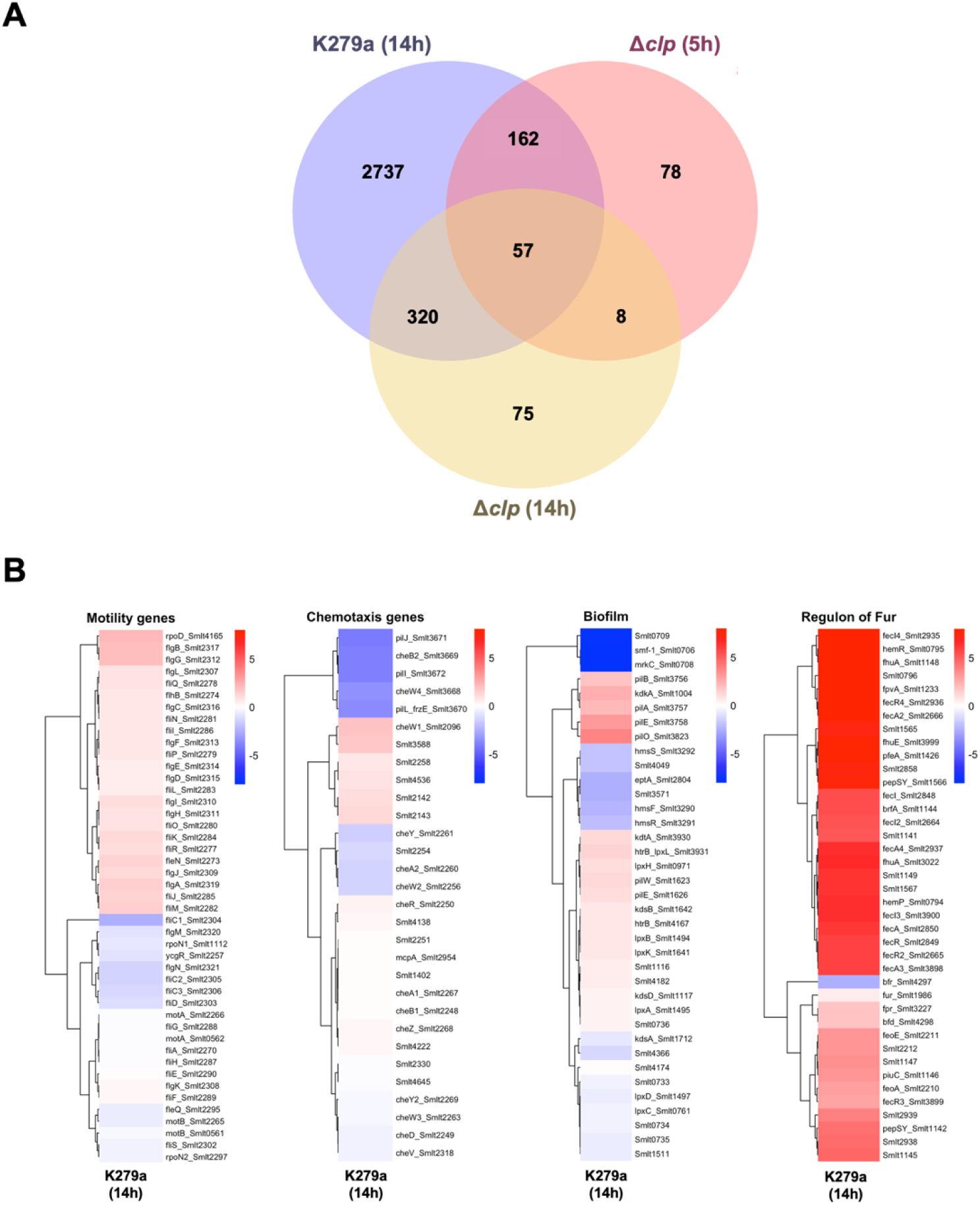
Growth phase–dependent regulation of Clp target genes in *S. maltophilia* K279a. (A) Venn diagram showing the overlap of differentially expressed genes (DEGs) identified in wild-type cells between 14 h and 5 h, in Δ*clp* at 5 h, and in Δ*clp* at 14 h, illustrating the growth-dependent regulatory role of Clp. (B) Heat maps of DEGs associated with motility, chemotaxis, biofilm formation, and iron homeostasis (Fur regulon) in the wild type at 14 h compared with 5 h. Red and blue indicate up- and down-regulation, respectively.

**S4 Figure.**
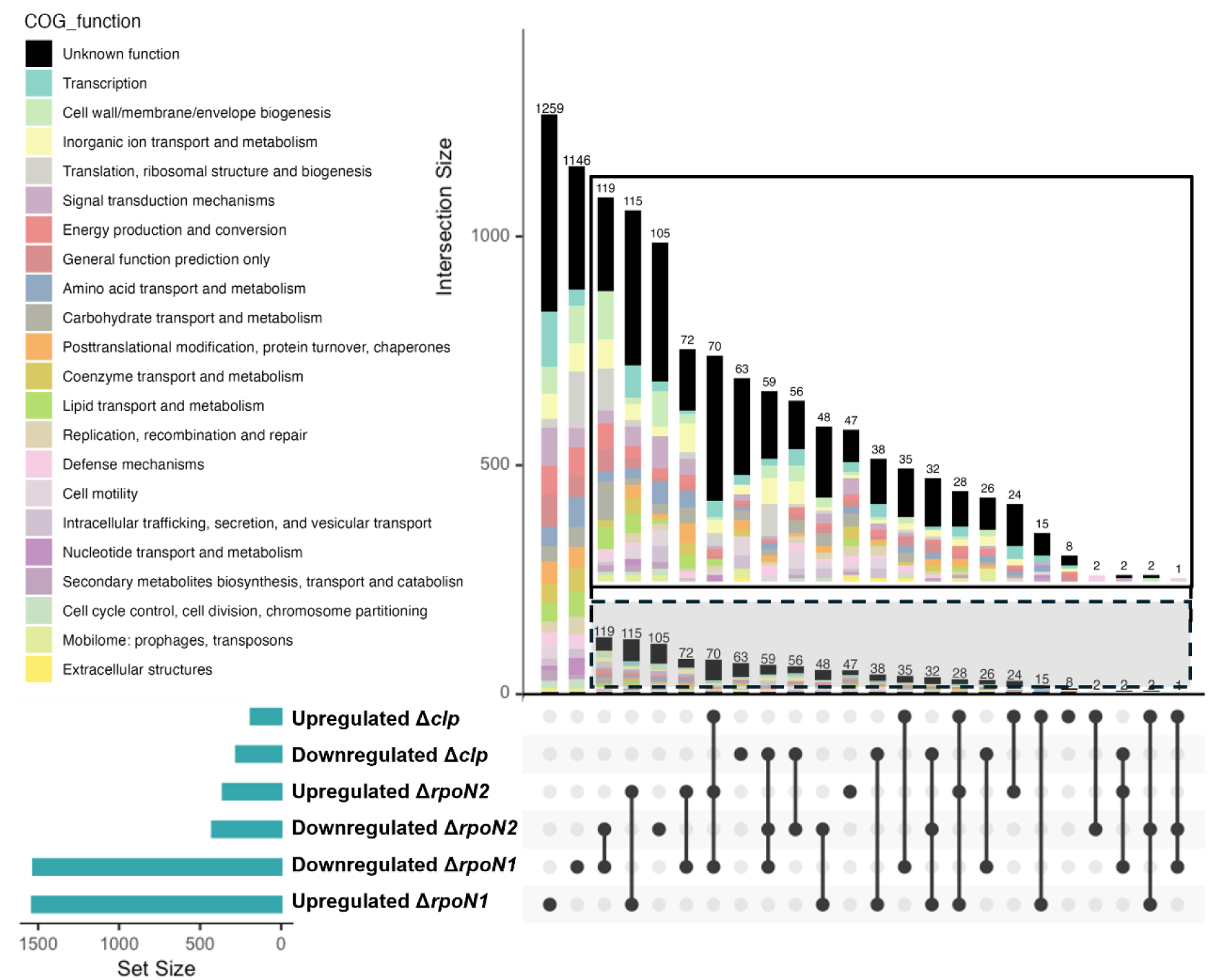
Shared transcriptional regulation between Clp, RpoN1, and RpoN2 in *S. maltophili*a K279a. UpSet plot showing the overlap of differentially expressed genes (DEGs) among Δ*rpoN1*, Δ*rpoN2*, and Δ*clp* mutants at 14 h (DSF quorum-sensing activation). Bars represent the size of intersecting DEG sets, which are further classified by COG functional categories (coloured). Set size indicates the number of DEGs identified in each individual regulon.

**S5 Figure.**
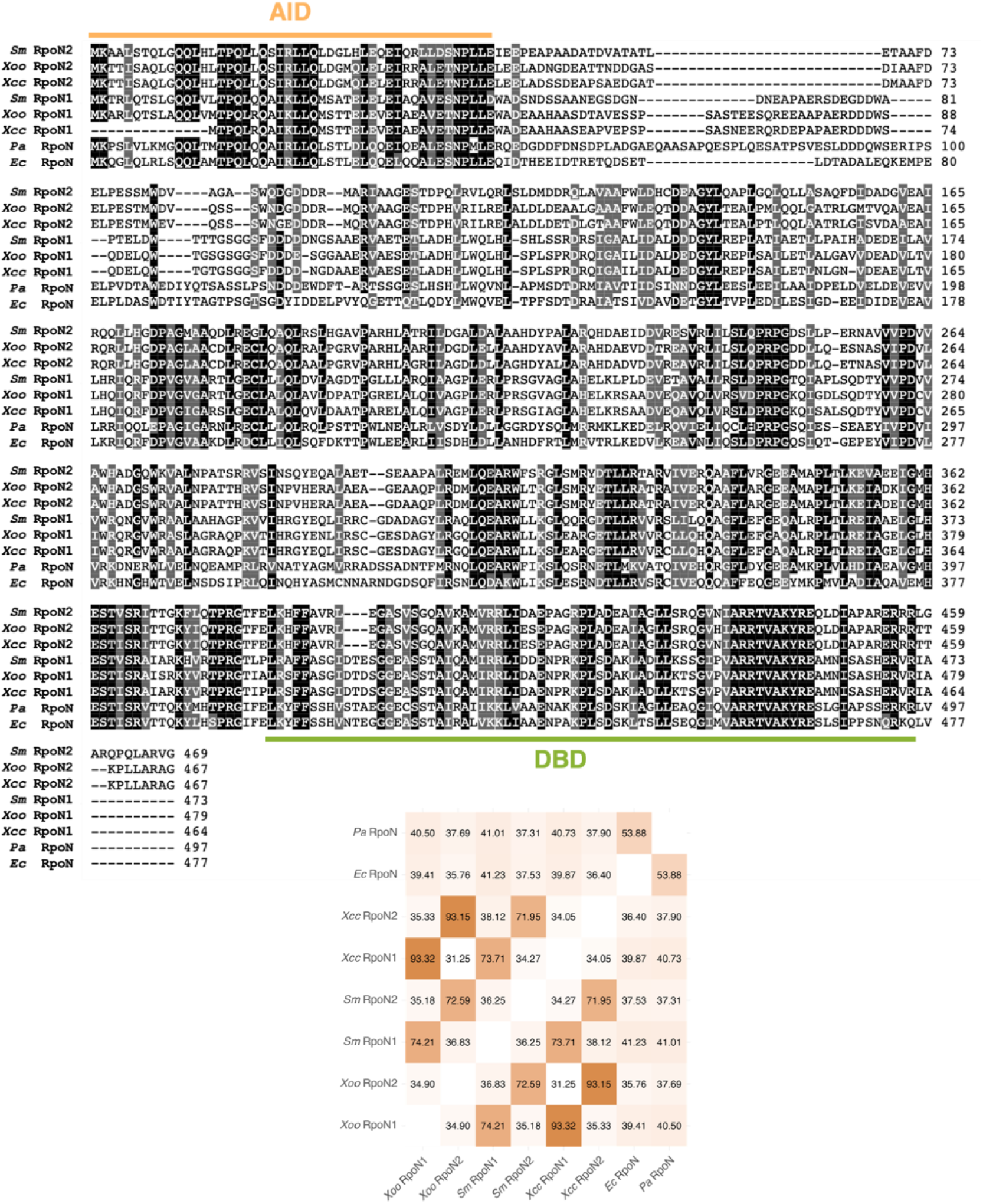
Conserved domains and sequence identity of σ⁵⁴ factors RpoN1 and RpoN2. Multiple sequence alignment of RpoN proteins from *S. maltophilia* (*Sm*), *X. campestris* (*Xcc*), *X. oryzae* (*Xoo*), *P. aeruginosa* (*Pa*), and *E. coli* (*Ec*). Conserved domains are indicated, including the activator interaction domain (AID, orange box) and the DNA-binding domain (DBD, green box). Alignment was performed using ClustalW, and gray shading indicates conserved amino acids among sequences. Bottom panel: pairwise identity matrix showing the percentage of identical residues among the RpoN proteins analyzed.

**S6 Figure.**
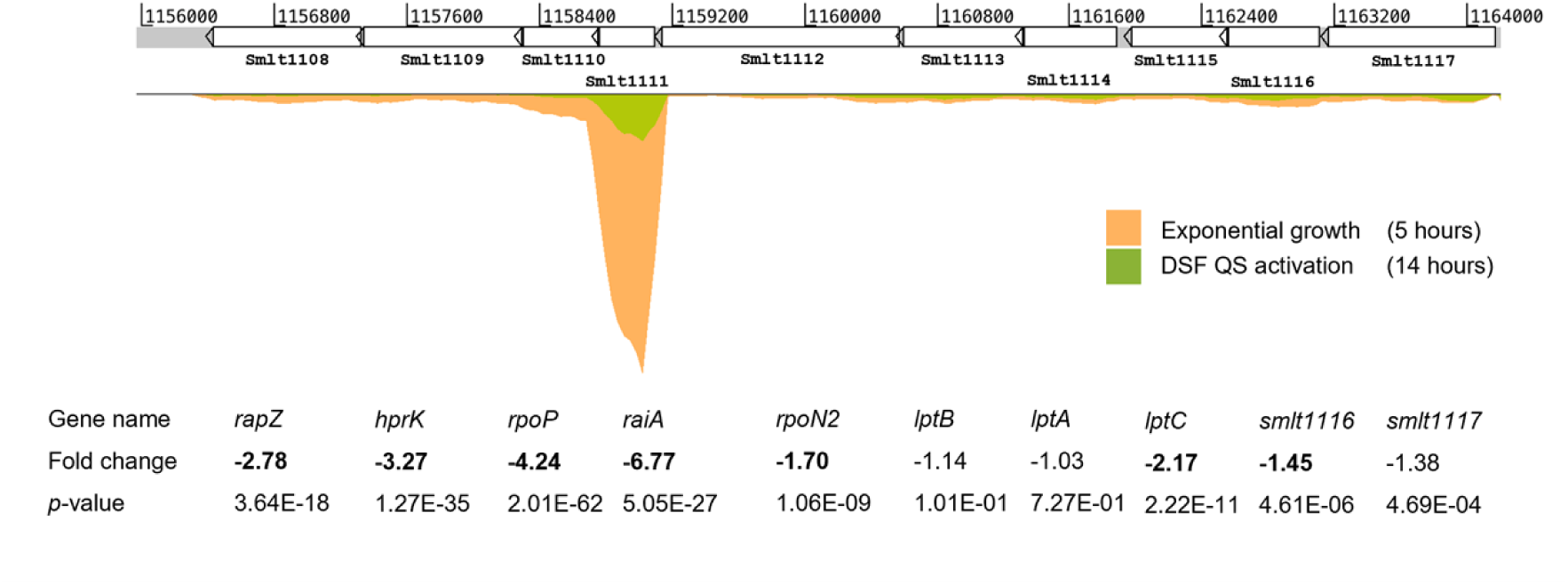
Growth phase–dependent downregulation of the rpoN1 operon at stationary phase entry. RNA-seq read coverage across the *rpoN1* operon (smlt1112 and associated genes) in *Stenotrophomonas maltophilia* K279a, comparing samples collected at 14 h (onset of stationary phase) with those at 5 h (exponential growth). Fold-change values indicate transcriptional variation between the two conditions; significant changes are highlighted in bold.

**S7 Figure.**
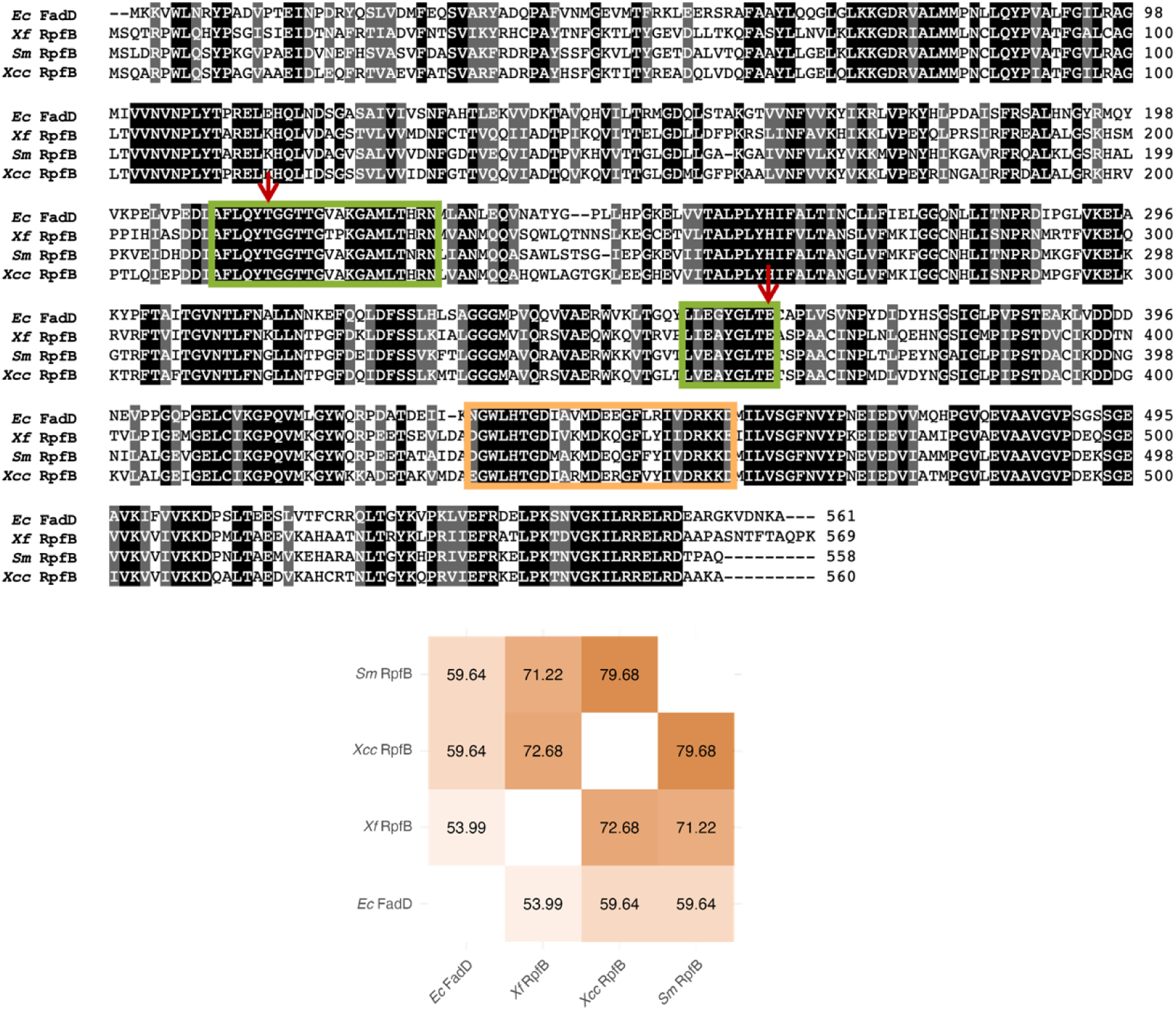
Conserved motifs and sequence similarity of RpfB proteins. Multiple sequence alignment of the long-chain fatty acid-CoA ligase (FCL) RpfB from *S. maltophilia* (*Sm*), *X. campestris* (*Xcc*), and *X. fastidiosa* (*Xf*), and the FadD protein from *E. coli* (*Ec*). The conserved ATP/AMP-binding motifs (green boxes) and FCL signature motif (orange box) are highlighted. Red arrows mark the conserved threonine and glutamate residues corresponding to the predicted active sites. Bottom panel: pairwise identity matrix showing the percentage of identical residues among the analyzed sequences.

**S1 Table: Susceptibility profile of *rpf* and *clp* mutants.**

**S2 Table: Susceptibility profile of *rpoN1* and *rpoN2* mutants and complemented strains.**

**S3 Table: Percentage of specific fatty acids relative to total cellular fatty acids in the indicated strains grown in the specified media.**

**S4 Table: Strains used in this work.**

**S5 Table: Plasmids used in this work.**

**S6 Table: Primers used in this work.**

**S1 File. Differentially expressed genes among all samples.**

## Notes

### Competing Interest Statement

The authors have declared no competing interest.

